# Nuclear DNA replication in *Leishmania major* relies on a single constitutive origin per chromosome supplemented by thousands of stochastic initiation events

**DOI:** 10.1101/2024.11.14.623610

**Authors:** Jeziel D. Damasceno, Gabriel L. A. Silva, Catarina A. Marques, Marija Krasilnikova, Craig Lapsley, Dario Beraldi, Richard McCulloch

**Affiliations:** The University of Glasgow Centre for Parasitology, The Wellcome Centre for Integrative Parasitology, University of Glasgow, School of Infection and Immunity, Sir Graeme Davies Building, 120 University Place, Glasgow, G12 8TA, United Kingdom

## Abstract

Understanding genome duplication requires characterisation of the locations where DNA replication initiates, termed origins. Genome-wide mapping of DNA replication origins has mainly been derived from population-based techniques, with only a few studies examining origin location and usage at the single-cell or single-molecule level. *Leishmania* are protozoan parasites where the first attempt to map DNA replication suggested the unprecedented use, for a eukaryote, of just a single origin per chromosome, while a subsequent approach suggested around 200-fold more origins. To reconcile these data and understand DNA replication dynamics in *Leishmania major*, we have applied DNAscent, a deep learning assay that uses long-read Nanopore sequencing to detect patterns of BrdU incorporation in individual DNA molecules, allowing the description of DNA replication fork movement and prediction of initiation and termination sites across the parasite genome. Our findings confirm the pre-eminence of a single locus of DNA replication initiation in each chromosome and reveal that this locus alone is constitutively activated in S-phase, with bidirectional forks emerging from discrete sites at the ends of multigene transcription units. DNAscent also reveals a much larger number of DNA replication initiation events that have not been detected in any previous mapping and are used stochastically, but whose abundance is greater as chromosome size increases. We show that each of these stochastic initiation sites localise to regions with high AT content, increased G-quadruplex levels and lower chromatin occupancy. In addition, we find markedly increased stochastic DNA replication initiation at sites with lower levels of nascent RNA transcripts. Finally, we show that all DNA replication initiation events result in mutagenesis. This work reveals a novel, bimodal strategy for DNA replication programming in *Leishmania* that drives genome transmission, replication timing and variation.

## Introduction

Propagation of life depends upon the successful duplication of an organism’s genome. In cellular organisms, DNA replication initiates at defined sites in the genome termed origins^1^, which in eukaryotes are designated by the binding of the Origin Recognition Complex (ORC)^2^. Unlike in bacteria and some archaea, where the whole genome is replicated from a single origin, each eukaryotic chromosome is normally replicated from multiple origins. This increase in origin number is associated with increased complexity in DNA replication organisation. With the exception of *Saccharomyces cerevisiae* and related yeasts^3^, origins in eukaryotes differ from those in prokaryotes in that they are not conserved DNA sequences, but are defined by sequence-independent genomic features. In addition, though origins are designated by ORC binding in the G1 phase of the cell cycle and then activated in S-phase, not all designated origins are activated. Moreover, there is a temporal order to the time in S-phase when origins are activated^4,5^ and, in the case of multicellular organisms, both the number and location of origins can vary during development and differentiation^1^. Due to such complexity, mapping DNA replication origins in eukaryotic cells is technically challenging and results from different techniques are often discordant^6^.

Most approaches to detect origins rely on mapping the DNA replication machinery or capturing replicative DNA synthesis in populations of cells, with the potential that only frequently used origins are detected and cell-to-cell heterogeneity is overlooked^1,6,7^. These limitations have be addressed by the recent development of both genome-wide single-molecule DNA replication mapping^8–11^ and single-cell DNA replication sequencing^12–14^ approaches, which have revealed previously undetected origins and informed on DNA replication timing. In *Saccharomyces cerevisiae*, two related approaches (DNAscent and Fork-seq/NanoForkSpeed)^8,9,15^ used Oxford Nanopore Technologies (ONT) sequencing to detect incorporation of the thymidine analogue 5-bromodeoxyuridine (BrdU) into newly replicated DNA, revealing that around 10-20% of DNA initiation events do not localise to previously reported origins^8,9^. Whether these newly predicted initiation sites correspond with a predicted 15% of MCM binding sites that do not localise with sequence-conserved origins^16^ awaits testing. In the larger genome of humans, ∼80% of origins detected by DNAscent do not overlap with origins previously detected by population-level analyses (Carrington et al BioRXiv 10.1101/2024.04.28.591325), perhaps consistent with a model of highly stochastic DNA replication initiation derived from optical replication mapping^10^. Single-cell DNA replication sequencing methods, such as scRepli-seq, measure copy number variation (CNV) between replicating and non-replicating DNA and, instead of mapping origin location, describe replication timing domains across S-phase^13,14^. These approaches have been used in human and mouse cells and, overall, reveal conservation of DNA replication timing organisation in individual cells and equivalent populations^13,14^, although timing domains differ between cell types^12,14^. Thus, despite origin activation being stochastic, the mammalian DNA replication timing program is spatially well defined, perhaps to ensure maximal efficiency of genome duplication and coordination with other processes, such as transcription.

Protozoans provide much of the diversity in the eukaryotic domain^17,18^ and only relatively recently has DNA replication been examined in a few select organisms in this grouping, with new insights into origin usage and DNA replication programming emerging. For instance, two nuclei are found in the single cells of *Giardia lamblia*^19^ and *Tetrahymena thermophila*^20^, with highly distinct genome organisation and function found in the distinct nuclei of the latter. In both cases, DNA replication dynamics appear to differ between the nuclei, but no work has mapped origins genome-wide, and so how potentially distinct genome duplication activities are organised within a single cell remains unclear. The life cycle of the malaria parasite *Plasmodium* is complex, containing four stages with distinct replicative strategies: hepatic and erythrocytic schizogony, gametogenesis, and sporogony^21^. Recently, two studies have combined ChIP-seq of ORC subunits and NanoForkSpeed or DNAscent to detect origins in *Plasmodium falciparum* undergoing erythrocytic schizogony^22,23^, a process in which the parasite undergoes multiple rounds of asynchronous DNA replication and nuclear division without cytokinesis, resulting in a multinucleated schizont.

Surprisingly, no clear consensus emerged from the two studies, perhaps suggesting that DNA replication during schizogony has unanticipated complexities. Arguably, the most advanced understanding of protozoan DNA replication has emerged within the kinetoplastids^24,25^, a ubiquitous grouping of flagellated organisms that includes both human and animal parasites of clinical and economic importance. DNA replication dynamics has been described and compared in three kinetoplastid parasites (see^26–31^ for reviews): *Trypanosoma brucei*^32–34^, *Trypanosoma cruzi*^35,36^ and *Leishmania* sp.^37–39^. Despite each of these parasites using a common, highly unusual form of gene expression where nearly every gene is expressed from a polycistronic transcription unit (PTU)^40^, and each genome sharing considerable synteny^41,42^, population-level DNA replication analyses suggest pronounced differences, in particular between *T. brucei* and *Leishmania*^31^.

The genome of *T. brucei* is primarily housed in 11 diploid ‘megabase’ chromosomes, each of which contains a highly transcribed core and two largely transcriptionally silent subtelomeres, which are variable in content between strains^43^ and mainly harbour thousands of genes encoding Variant Surface Glycoproteins (VSGs)^44–46^. *T. brucei* DNA replication has to date only been mapped genome-wide using population-level Marker Frequency Analysis sequencing (MFA-seq; equivalent to sort-seq^47^ in yeast)^32–34,48,49^. This approach predicts origins within the megabase chromosome cores at the boundaries (‘strand switch regions’, SSRs) of ∼25% of PTUs. Consistent with these loci being origins, one subunit of *T. brucei* ORC^50–52^ has been shown to bind to all SSRs, suggesting that only a subset of ORC-binding sites are activated to initiate DNA replication in S-phase^32^, with such origin selection being invariant between distinct life cycle stages and parasite strains^33^. Amongst these MFA-seq-predicted origins, those co-localising with centromeres are the earliest replicating^32,33^. More recent MFA-seq mapping suggests that most of the subtelomeric compartment of the *T. brucei* genome is late-replicating and more unstable than the core^34^. Altogether, these data reveal incompletely explored links between DNA replication initiation, transcription, chromosome segregation and genome stability in this organism^32–34,48,49^.

Equivalent MFA-seq analysis in two *Leishmania* species, *L. major* (36 chromosomes) and *L. mexicana* (34 chromosomes), revealed a striking difference to *T. brucei*: only a single MFA-seq peak indicative of S-phase DNA replication initiation could be detected in each chromosome^53^. As no study has mapped ORC in either *Leishmania* genome, it is premature to say that the MFA-seq signal in each chromosome represents an origin, but a range of observations are consistent with such a suggestion: first, as in *T. brucei*, each MFA-seq signal centres on an SSR; second, mapping the binding of the kinetochore subunit KKT1 is consistent with the single MFA-seq SSR in each chromosome being a centromere^54^, which are coincident with the earliest acting origins in *T. brucei* ^32^; third, ∼40% of the *Leishmania* MFA-seq SSRs are syntenic with origin-active SSRs in *T. brucei*^53^; and, finally, the single MFA-seq peak in each chromosome overlaps with highly localised changes in base composition skews^38^ as well as with increased mutagenesis^39^, each feature being consistent with sites of frequent DNA replication initiation. Beyond these data on the putative *Leishmania* origin-active SSRs themselves, average chromosome replication timing in *L. major* is unusual in that it correlates with chromosome length, with larger chromosomes being replicated later than smaller ones^38^. Such timing may be consistent with limiting DNA replication initiation to just a single locus in each chromosome, but such programming appears insufficient to allow duplication of the largest ∼40% of chromosomes during S-phase^53^. At least in part, this limitation may be overcome by DNA replication activity proximal to the telomeres of each *L. major* chromosome that is detectable outside S-phase^38^. Nonetheless, two different approaches have suggested that MFA-seq may not detect all DNA replication activity in *Leishmania*. DNA combing has detected >1 site of DNA replication initiation in single *Leishmania* DNA molecules^55,56^, and Short Nascent Strand-sequencing (SNS-seq) has mapped >5000 putative origins throughout the *L. major* genome, with only limited overlap with MFA-seq mapping^57^. These discrepancies may arise from technical limitations in each of these methodologies. MFA-seq analyses CNV in replicating versus non-replicating cells, and therefore mainly determines the timing of DNA replication, with limited spatial resolution to pinpoint origins. DNA combing as reported to date in *Leishmania* is low-throughput and does not localise where in a chromosome, or even in what genome, DNA replication events are detected; indeed, it cannot exclude the possibility that the multi-origin molecules described are derived from abundant extrachromosomal elements^58^. Finally, SNS-seq mapping is known to be susceptible to G-quadruplex (G4) impediments^59^, which may be a particular concern given the high prevalence of G4s in the *L. major* genome^60^.

Irrespective of technical considerations, the huge discrepancies between these studies in estimates of origin number and location raise questions about how *Leishmania* DNA replication is programmed, including why MFA-seq portrays such a stark difference relative to *T. brucei*. Here, we address these questions using DNAscent, a deep learning-based approach developed by Muller and colleagues^8,61^ that, by detecting BrdU in newly replicated DNA on ultra-long, single-molecule Nanopore sequence reads, provides high-precision mapping of DNA replication fork dynamics, allowing the detection of sites of DNA replication initiation, termination and pausing. Using this approach, we confirm that the single MFA-seq signal in each *L. major* chromosome represents the pre-eminent locus at which DNA replication initiates at the start of S-phase, and that such constitutive, cell cycle-dependent activity is supported by much more numerous and stochastic sites of DNA replication initiation. We also provide evidence that chromosome length-related DNA replication timing in *L. major* is reflected in differential density of the stochastic initiation events among chromosomes throughout S-phase, and that these events localise to regions with high AT content, increased G-quadruplex levels and lower chromatin occupancy. This imbalance between constitutive and stochastic DNA replication initiation appears extreme amongst eukaryotes described to data, and we provide evidence that such an unusual bimodal DNA replication programme relates to patterns of genome variation in *L. major*.

## Results

### BrdU detection with DNAscent confirms pre-eminent DNA replication initiation at a single locus in each *L. major* chromosome

DNAscent relies on the detection of BrdU in DNA molecules through nucleotide analogue signal currents during Nanopore sequencing (Fig.1A). To establish that detection of BrdU incorporated into the nuclear genome *Leishmania* is possible (for instance, it is not impeded by signal from the hypermodified thymidine base, β-D-Glucopyranosyloxymethyluracil, also called base J)^62^, *L. major* promastigote (insect stage) cells were arrested at the G1/S-phase transition of the cell cycle by treatment with 5 mM hydroxyurea (HU) for 8 hours. Next, one sample was collected (no BrdU control), while the remainder of the cells were released from the HU block and grown for 15, 30 or 60 min in the presence of 150 µM BrdU (Fig.S1A). This treatment was followed by a thymidine-chase (1 mM thymidine, 60 min), DNA extraction, DNA sequencing on a ONT MinION, and BrdU calling with DNAscentv2^8,61^. Barely any predicted BrdU signal was seen in the no BrdU control cells (0.17% of reads), while substantial BrdU signal accumulated, in a time dependent manner, in the BrdU labelled cells (1.33-2.24% of reads). Moreover, metaplots (Fig.S1B) showed that most BrdU signal accumulated around the single SSR previously predicted by MFA-seq to be the single early replicating locus in each chromosome^38,53^.

**Figure 1.**
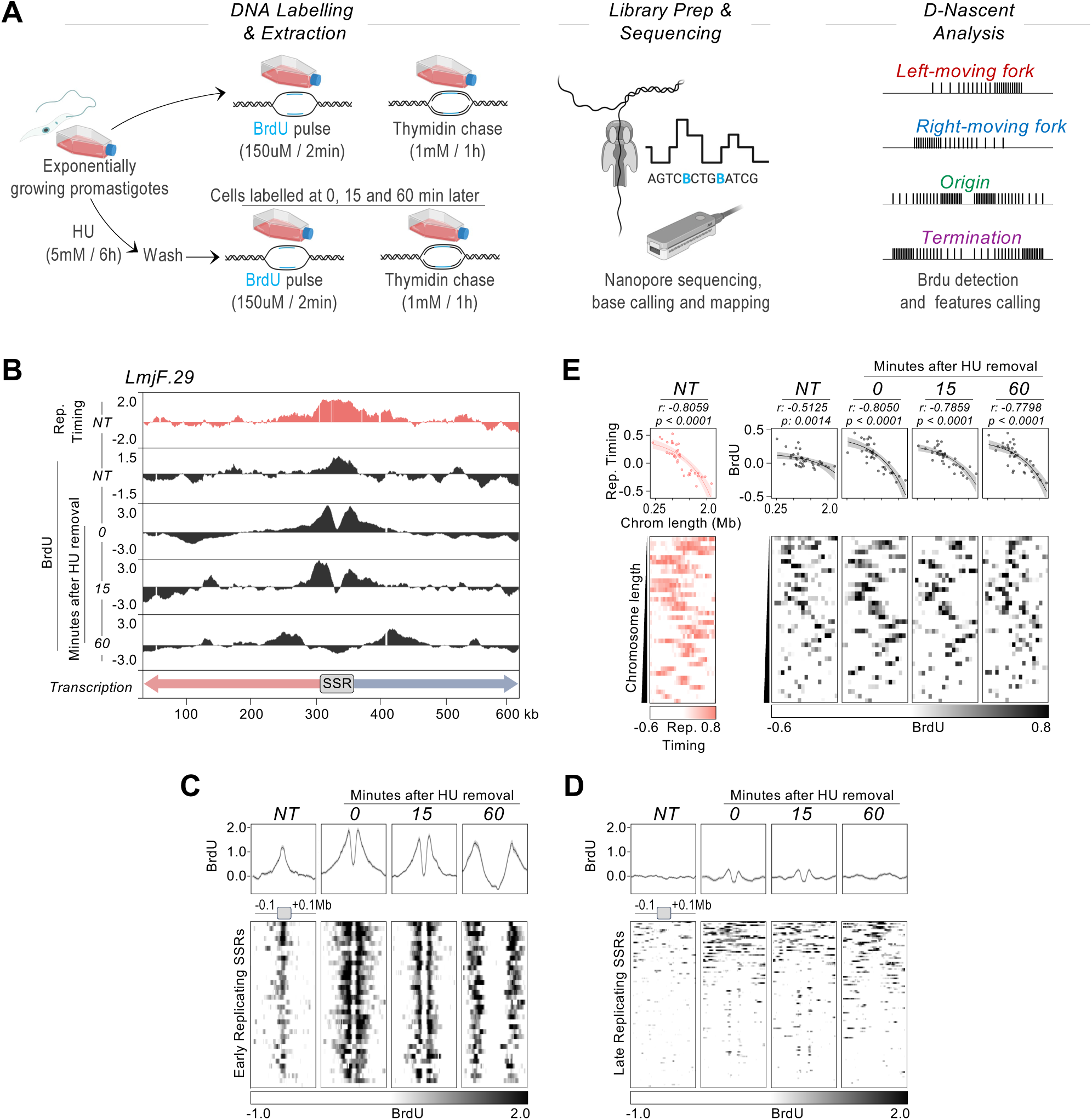
Mapping BrdU incorporation profiles with DNAscent confirms centromeric SSR-driven DNA replication initiation and chromosome size-associated DNA replication timing. A) Schematic of the experimental approach. Exponentially growing *L. major* promastigotes (NT) were labelled with BrdU for 2 minutes followed by a thymidine chase for 1 hour. The same labelling conditions were also used 0, 15 and 60 minutes after parasites were released from cell cycle arrest at G1/S with HU. High molecular weight DNA was extracted and subjected to Oxford Nanopore Technologies sequencing, and the data then analysed with DNAscentv2. B) Snapshot showing BrdU scores (black) around a representative early-replicating SSR in *L. major* chromosome 29 in NT cells and 0, 15 and 60 minutes after HU removal. The top track (salmon) shows the DNA replication profile in the same region as determined by MFA-seq. C) and D) Metaplots showing BrdU score profiles around all early-replicating SSRs and all late-replicating SSRs, respectively, in NT and HU synchronised cells. E) Colourmaps (lower panels) show average replication timing of each chromosome determined by MFA-seq (left, salmon), and average BrdU score per chromosome (right, black; NT cells, and 0, 15 and 60 minutes after release from HU arrest are shown). Top panels show linear regression between chromosome length and average MFA-seq timing or average BrdU scores; r and p values are indicated at the top of each panel and shading represents 95% confidence intervals.

To accurately predict replication fork movement, as well as initiation and termination sites of DNA replication, DNAscent relies on the detection of gradients of BrdU incorporation (Fig.1A)^8,61^. We therefore modified the above experiments, again with *L. major* promastigotes, but this time reducing the length of the BrdU pulse to 2 mins: since the rate of replication fork movement in *L. major* has been calculated as ∼2.5-2.8 kb/min ^55,57^, we expected BrdU labelled tracts of ∼5-5.6 kb, which is smaller than the N50 of 20.5 kb we recover from Nanopore sequencing. For this new experimental setup, we again arrested the cells with HU at the G1/S-phase transition and labelled with 150 µM BrdU at 0 min, 15 min and 60 min after release. In addition, we labelled cells with 150 µM BrdU for 2 mins without any HU treatment, allowing us to compare patterns of BrdU incorporation, and thereby DNA replication dynamics, in the absence of HU-mediated cell cycle synchronisation relative to the early-mid stages of S-phase after HU stall release (Fig. 1A). (In all cases, the BrdU pulse was followed by a 1 mM thymidine chase of 60 min.)

Visual inspection revealed that BrdU signal enrichment across the chromosomes matched previous MFA-seq analysis, with the most prominent level of BrdU in unsynchronised cells found at a single locus in each chromosome that overlapped the SSR on which MFA-seq signal accumulates (Fig.1B; Fig.S2). After HU release, this most prominent BrdU signal was detected as two peaks of increasing separation with time, consistent with bidirectional progression of DNA replication from the SSR (Fig.1B; Fig.S2). To test these observations further, we generated metaplots of BrdU signal around the single SSR in each chromosome that displayed an MFA-seq peak (36 ‘early-replicating SSRs’), and around all other SSRs (Fig.1C, D; ‘late-replicating SSRs’). Very consistent levels of BrdU signal were apparent at the MFA-seq enriched, early-replicating SSR in each chromosome both before and after HU synchronisation: whereas BrdU signal was closely focused around the SSR in unsynchronised cells, two BrdU peaks of very similar amplitude and width were detected flanking the SSRs in the HU-synchronised cells, with the distance between the peaks increasing over time after release from HU arrest (most distal in the 60 min sample). Taken together, these data indicate highly coordinated initiation of DNA replication in S-phase at a single SSR in each chromosome (Fig.1C). In contrast, there was no such clear BrdU signal enrichment at the other, late-replicating SSRs in the genome, with or without HU synchronisation (Fig.1D).

Using MFA-seq data, we have previously noted a DNA replication programme in *L. major* where average chromosome duplication timing is size-dependent (Fig. 1E, left panel, coloured salmon)^27,38,53,63^. To test if DNAscent detects the same organisation of DNA replication, linear regression analysis was conducted on the BrdU density data, which revealed a significant correlation between chromosome size and average BrdU signal, with smaller chromosomes displaying higher BrdU levels than larger chromosomes in non-synchronized cells (Fig. 1E, panel NT, coloured black). An even stronger correlation was observed in cells after release from G1/S synchronisation, demonstrating that the earlier replication of smaller chromosomes occurs across S-phase (Fig. 1E, panels 0, 15 and 60, coloured black). Importantly, this effect cannot be attributed to differing levels of Nanopore read depth across the chromosomes (Fig.S3A), and linear regression analysis showed there was no equivalent size-dependent correlation between chromosome length and base T content (Fig.S3B), ruling out BrdU distribution being due to skews in sequencing coverage or base content.

Taken together, these data show that DNAscent analysis of long-read Nanopore sequencing of BrdU-labelled DNA is not only feasible in *L. major* but validates the findings of population-level MFA-seq mapping: in early S-phase there is highly coordinated initiation of DNA replication from a single locus at each chromosome, which progresses bidirectionally towards chromosomes ends, and which may dictate chromosome size-dependent replication timing.

### Origin prediction with DNAscent reveals abundant, previously undetected initiation events in each *L. major* chromosome

The presence of a single, early replicating locus on each *L. major* chromosome, as first indicated by MFA-seq^38,53^ and here confirmed by BrdU density in DNAscent (Fig.1), suggests two potential scenarios of genome duplication: either each chromosome is replicated from a single putative origin (or a single cluster of origins), or each chromosome is replicated by multiple origins that fall below the detection limits of MFA-seq and may be activated at different stages of S-phase. An advantage of DNAscent^8,61^ or variants^9,11,15,64^ over population-level mapping approaches lies in their higher sensitivity, as they can predict DNA replication initiation and termination sites, based on fork direction determined from BrdU gradients (Fig.1A) on single DNA molecules that collectively span the genome. Thus, heterogenous aspects of DNA replication that are lost in population-based methodologies as signals are averaged across many cells and molecules may be revealed. Visual inspection of individual Nanopore ultra-long reads after processing with DNAscent revealed DNA molecules containing multiple sites of DNA replication initiation (Fig. 2A).

**Figure 2.**
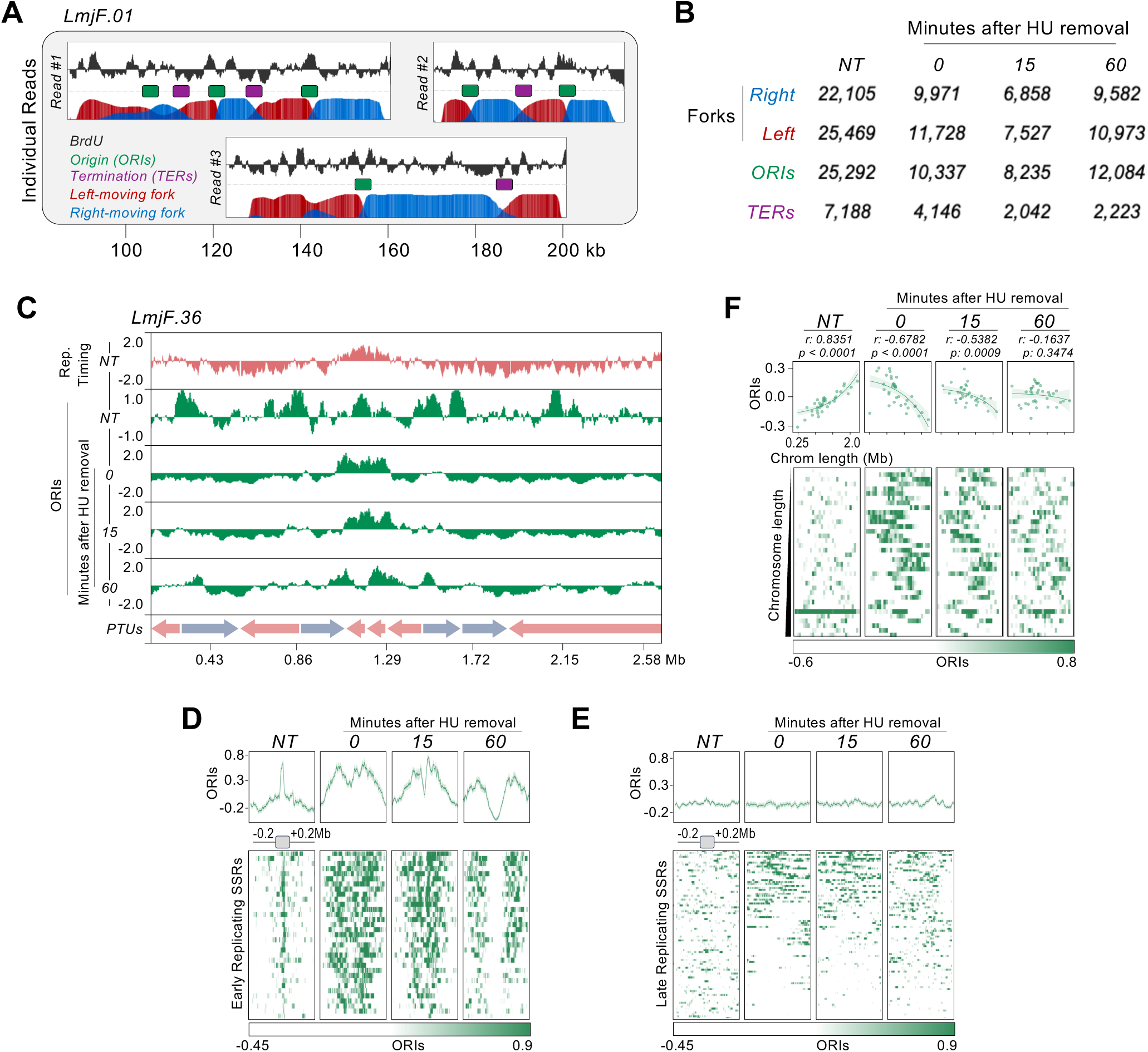
DNAscent shows that a single centromeric SSR in each chromosome is activated in early S phase and reveals widespread initiation events whose distribution is chromosome size-dependent. **A)** Representative Nanopore reads showing BrdU scores, probabilities for right- and left-moving DNA replication forks, and predicted DNA replication initiation (putative origins, ORIs) and termination (TER) sites. **B)** Summary of the numbers of DNA replication features predicted by DNAscent in unsynchronised (NT) cells and in cells 0, 15 and 60 mins after release from HU cell cycle stall. **C)** Snapshot showing ORI density (green) around a representative early-replicating SSR in NT cells and 0, 15 and 60 mins after release from HU stall. The top track (salmon) shows the DNA replication profile as determined by MFA-seq. **D)** and **E)** Metaplots showing global ORI density profile around all early-replicating SSRs and all late-replicating SSRs, respectively, both in NT and HU synchronised cells. **F)** Colourmaps (lower panels) show average ORI density per chromosome in NT cells and 0, 15 and 60 mins after release HU cell cycle arrest; top panels show linear regression between chromosome length and average ORI density; r and p values are indicated at the top of each panel, and shading represents 95% confidence intervals.

Moreover, comparison of reads derived from common regions of a chromosome suggest considerable variation in the locations of initiation sites (Fig. 2A). By examining all reads obtained, DNAscent predicted ∼25,000 initiation sites in non-synchronised cells and ∼8,200-12,100 initiation sites in HU-synchronised cells (Fig.2B). This difference is likely explained by the data from each HU synchronised sample capturing cells at a limited part of S-phase, whereas the non-synchronised sample consists of cells in all stages of S-phase (and indeed of the cell cycle). However, these predictions will overestimate of the total number of DNA replication initiation sites, as many reads overlap and therefore some sites will be overcounted as they are common to the overlapping reads. Nonetheless, DNAscent suggests a density of DNA replication initiation events substantially greater than inferences made from DNA combing^55^ (∼168 per haploid genome) and even from SNS-seq mapping^56^, which predicts ∼5,100. Such a conclusion is consistent with analysis of the distance between DNAscent-predicted initiation sites (inter ‘origin’ distance; IOD) on individual reads: a median DNAscent IOD of ∼20.7-22.3 kb (Fig. S4; range 0.48-160 kb) is at least 3-4 fold closer than ∼72-193 kb IODs reported by DNA combing^55,57^.

### ORIs predicted by DNAscent are non-randomly distributed across the *L. major* genome

The above data predict the existence of numerous, hitherto undetected DNA replication initiation events. As we do not know if any or all these predicted initiation sites correspond with ORC-defined origins, we will refer to them as ORIs. To capture the genome-wide distribution of DNAscent ORIs, we first determined aggregated (Fig.S5) and relative ORI density (Fig.2C, Fig.S5), allowing us to compare ORI distribution with MFA-seq mapping in unsynchronised and HU-synchronised cells (Fig.2C, Fig.S5). Unlike the clear correspondence between BrdU density and MFA-seq signal (Fig.1B, Fig.S2), visual inspection suggested a wider distribution of ORIs than MFA-seq signal across chromosomes in unsynchronised cells (Fig.2C, Fig.S5; HU non-treated, NT). Nonetheless, metaplots showed that the single early-replicating SSR at which MFA-seq signal accumulates in each chromosome was a locus of very marked, localised ORI accumulation (Fig.2D). In contrast, no such ORI localisation was seen at all other, late-replicating SSRs in the absence of HU synchronisation (Fig.2E), meaning other chromosome regions with higher relative ORI density to do not predominantly coincide with locations where transcription starts or ends. When we plotted ORI density across the genome in unsynchronised cells we found a significant correlation with chromosome size, with greater density of ORIs seen in larger chromosomes (Fig.2F), the opposite of the chromosome size-dependent distribution of DNA replication activity detected by both MFA-seq^38^ and BrdU density (Fig.1E).

HU-mediated cell cycle arrest and release substantially altered the patterns of ORI distribution. Chromosome-wide visualisation (Fig.2C, Fig.S5) revealed that ORI density was no longer distributed across the chromosomes after HU treatment, but instead largely colocalised with the MFA-seq signal immediately after HU exposure and spread outwards from these early-replicating SSRs 15 and 60 mins after release from HU. Metaplot analysis (Fig.2D) confirmed this: a pronounced accumulation of DNAscent ORIs was seen around the 36 MFA-seq, early-replicating SSRs immediately after HU release, with areas of high ORI density seen extending to both flanks of the SSRs in the 15- and 60-min samples, an effect highly comparable to BrdU density mapping (Fig.1B,1C). In no HU-synchronised sample was there evidence for ORI accumulation at other SSRs (Fig.2E). These data suggest that duplication of each *L. major* chromosome starts with DNA replication mainly focused on a single SSR in early S-phase. To test this interpretation, we performed linear regression analysis to compare average ORI density and chromosome length in the HU-synchronised samples (Fig.2F). This analysis showed that chromosome size-dependent ORI density was reversed by HU treatment relative to unsynchronised cell: whereas ORI density was greatest in the larger chromosomes in untreated cells, immediately after HU release ORI density was markedly greater on the smaller chromosomes compared with the larger chromosomes (Fig.2F). In addition, this distribution was less clear 15 mins after HU release, and no size-dependent ORI density was seen after 60 mins.

Taken together, the above data reveal a number of features of DNA replication programming in *L. major*. First, DNA replication is predominantly activated at a single SSR in each chromosome as cells enter S-phase. Second, this cell cycle-regulated, and likely constitutive activity is supported by much wider, stochastic DNA replication initiation events (ORIs) that are more common as chromosomes increase in size and may be more prevalent later in S-phase, suggesting this activity is needed to allow the complete duplication of the larger chromosomes. Finally, we see no evidence in *L. major* promastigotes for the stochastic ORIs only localising to SSRs, meaning the abundant DNA replication initiation events we now detect may not be spatially limited by polycistronic transcription, suggesting they differ from ORC-defined origins in *T. brucei* ^32^.

### Origin Efficiency Metrics (OEM) analysis provides a genome-wide mapping of DNA replication initiation and termination

One of the major advantages of DNAscent over MFA-seq, SNS-seq and DNA combing is that it can detect and map individual leftward and rightward moving DNA replication forks. By aggregating the DNAscent-mapped replication forks, similar to analysis performed with FORK-seq in yeast ^9,15,64^, we were able to generate genome-wide profiles of Replication Fork Directionality (RFD) as well as Origin Efficiency Metrics (OEM)^65–67^. RFD profiling provides a view of the predominant direction of replication forks at any given locus, with RFDs of −1.0 and 1.0 indicating 100% of left-or right-moving forks, respectively. OEM profiling indicates the extent of the upwards or downwards shifts in RFD profile, with positive and negative OEM values indicating sites of divergent (initiation zones, IZs) and convergent (termination zones, TZs) replication forks, respectively. OEM values of +1.0 and −1.0 indicate, respectively, an IZ or a TZ predicted to be constitutively activate in 100% of cells within the population.

Fig.3A shows RFD and OEM profiles in unsynchronised cells focused on the single, early-replicating SSR (as predicted by MFA-seq and confirmed by DNAscent) on *L. major* chromosome 4; this SSR is a site where the surrounding multigenic transcription units converge (ie it is a region of transcription termination). Fig.S6 shows further early-replicating SSRs where transcription units diverge (bi-directional initiation), or where surrounding multigenic transcription initiates and terminates. In all cases, the shift in RFD was notably localised within the SSR. In addition, all OEMs were notably high (Fig.3A, Fig.S6), indicating DNA replication activation in most Nanopore reads. Thus, RFD and OEM metrics derived from DNAscent at early-replicating *L. major* SSRs previously detected by MFA-seq indicate constitutive activation of DNA replication at these sites in the population and suggests initiation at a relatively localised region of the wider SSR, consistent with focused BrdU density (Fig.1C) and ORI localisation (Fig.2D) in the absence of HU treatment.

**Figure 3.**
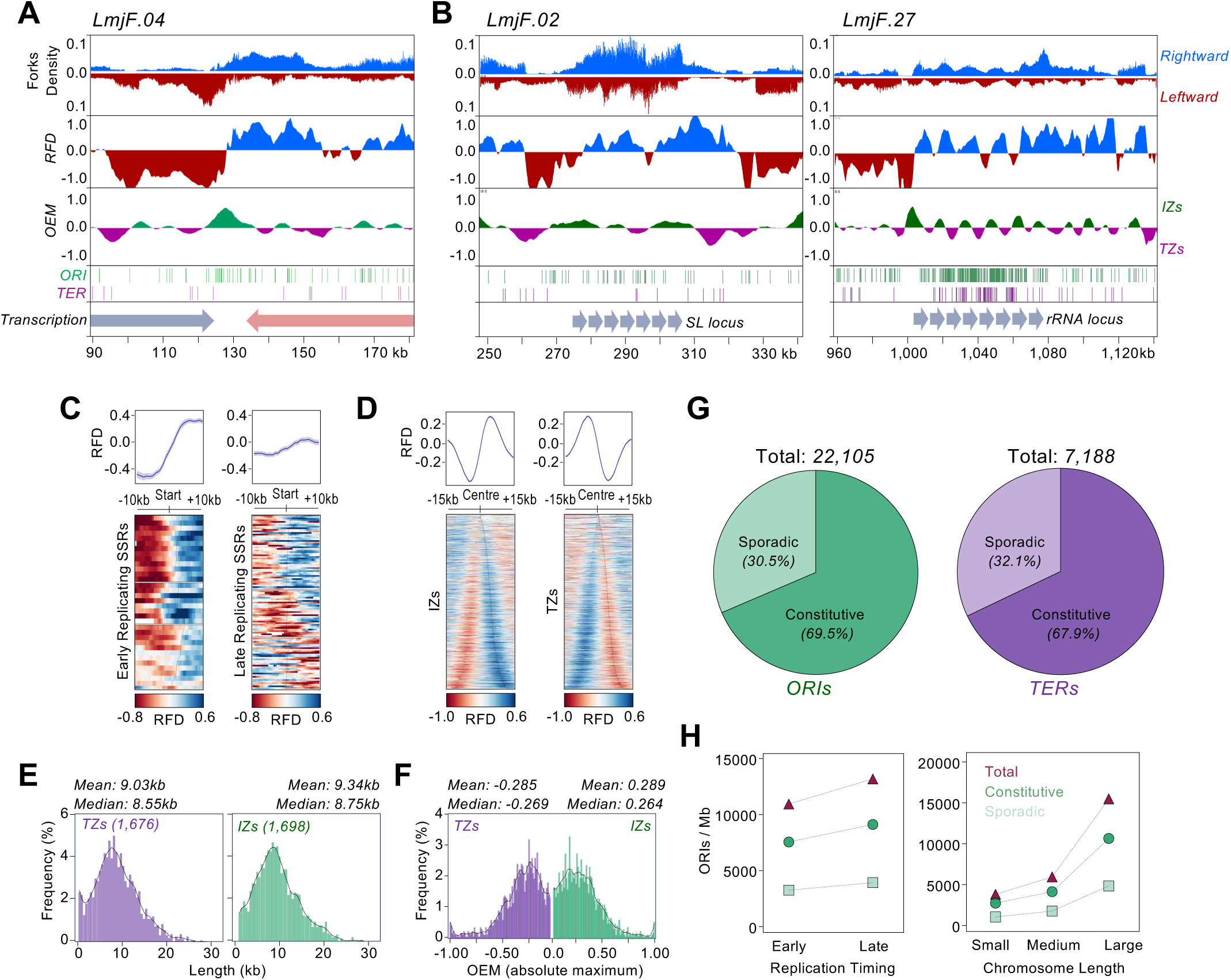
Analysis of DNA replication forks detected by DNAscent indicates genome duplication in *L. major* relies on stochastic DNA replication initiation. A) and **B)** Snapshots of three chromosome regions surrounding early-replicating SSRs, showing DNA replication fork density and quantification of predicted replication fork direction (RFD) and origin efficiency metrics (OEM); positive and negative OEM values indicate initiation (IZs) and termination (TZs) zones, respectively. **C)** Metaplots showing global RFD profiles around all early-replicating and all late-replicating SSRs in unsynchronised (NT) cells. **D)** Metaplots showing global RFD profiles around all IZs and TZs in NT cells. **E)** Global frequency distribution of IZ and TZ lengths. **F)** Global frequency distribution of OEM positive (IZs) and negative (TZs) values. **G)** Genome-wide quantification of constitutive and stochastic (‘sporadic’^9^) ORIs and TERs; constitutive ORIs and TERs were defined as those overlapping IZs and TZs, respectively, while ‘sporadic’ ORIs and TERs were defined as those that do not overlap TZs and IZs, respectively. **H)** Distribution of ORI types between the indicated genomic compartments.

Fig.3B shows the same analysis at two distinct SSRs, also predicted to be constitutively activated in early S-phase: here, MFA-seq predicted DNA replication to initiate at the boundaries of the splice-leader (chromosome 2) and rRNA (chromosome 27) loci^53^, each composed of repeated sequences. At the rDNA loci, a pronounced and localised RFD shift and high OEM value was seen around the predicted transcription start site (Fig.3B). However, the rightward fork (moving in the same direction as transcription) did not appear to move continuously across the locus, perhaps suggesting some DNA replication initiation at each repeat or pauses in fork progression, as has been seen in yeast^8^. The same high OEM value was not so clearly seen at the start of the splice leader locus, but instead a number of lower, positive OEMs were seen, perhaps suggesting multiple sites of DNA replication initiation across the repeats.

To examine the relationship between DNA replication initiation and RFD switching at a genome-wide level, we first performed metaplot analysis of the data around all SSRs (Fig.3C, Fig.S7A). This analysis revealed that RFD switching around all early-replicating SSRs, previously detected by MFA-seq, was significantly more pronounced than at all other, late-replicating SSRs (Fig. 3C). Furthermore, when comparing RFD switching between SSRs grouped according to the arrangement of flanking polycistronic transcription, we observed that RFD switching was more pronounced the higher the proportion of early-replicating SSRs in each group (Fig. S7A), which was reflected in higher OEM values (Fig.S7B). Taken together, these data suggest that sharper RFD switching at *L. major* SSRs is not explained by transcription. Thus, early replication timing, as initially predicted by MFA-seq, is a feature of specific SSRs, rather than being a means to accommodate the unusual nature of gene expression in *Leishmania* at all SSRs.

The above data is focused on the SSRs, where origins are detected in *T. brucei* ^26,32^. However, large number of regions of upwards and downwards RFD switching are seen across all the chromosomes (Fig. S6) and are not limited to SSRs, consistent with previously undetected, genome-wide initiation and termination events (Fig.1, Fig.2). Here, these loci should be considered as IZs and TZs, given that fork direction has been aggregated across Nanopore molecules (Fig. S6). OEM analysis revealed the existence of 1,698 IZs and 1,676 TZs, with an average length of 9.34 kb and 9.06 kb, respectively, suggesting DNA replication initiates approximately every 18.8 Kb (Fig 3E). Metaplot analysis confirmed the expected converging and diverging RFD profiles around IZs and TZs, respectively, as well as heterogeneity in the location of RFDs across fixed-size genome fragments (Fig. 3D). To examine how frequently these zones are found in the population, we examined the range of OEM values for both IZs and TZs (Fig.3F). Only a small fraction (∼0.5%) of IZs and TZs had an OEM close to +1.0 or −1.0, respectively, indicating very few of these regions are used in all cells (Fig. 3F). Instead, the mean OEM values for IZs and TZs were +0.289 and −0.285, respectively, indicating that the majority of these regions are found in only around 28% of cells, consistent with stochastic usage.

Calculating how many DNAscent-predicted ORIs and termination sites (TERs) in individual reads colocalise with the IZs and TZs identified by RFD switching and OEM provides a rough measure of whether a given ORI or TER is used constitutively or ‘sporadically’ in the population^9^. Here, ∼30% of ORIs and ∼32% of TERs predicted by DNAscent were found not to overlap with TZs and IZs, respectively, suggesting a high proportion of stochastic initiation and termination events (Fig.3G), substantially exceeding the 9% of sporadic origins predicted in yeast by a comparable analysis^9^.

Calculating the density (ORIs/Mb) of predicted constitutive and sporadic ORIs in early or late compartments of the chromosomes, as well as in chromosomes categorised by size, showed that both constitutive and sporadic origins were found more frequently in late-replicating regions of the genome (Fig.3H). These data are consistent with our previous demonstration of a higher density of DNAscent-predicted ORIs in the larger, later-replicating chromosomes (Fig.2F).

### IZs and TZs identified by DNascent are regions of localised variation in sequence content and chromatin accessibility

In an attempt to identify any features associated with the abundant, DNAscent-predicted stochastic DNA replication and termination sites, we performed metaplot analyses of all the IZs and TZs identified by OEM and examined their average sequence content (Fig.4A). Although no conserved sequences were found, IZs were notable as regions of increased AT content in the genome, whereas TZs were found in GC-enriched loci. In addition, there was some evidence of G4 enrichment on both DNA strands at IZs, whereas G4s were underrepresented at TZs. Finally, mapping of MNase-seq^57^ data showed that IZs were areas of low nucleosome occupancy, while TZs were enriched in nucleosomes. Each of these genomic features of IZs become more pronounced in more frequently used IZs (Fig.4B): when IZs were separated into 4 different groups according to their average OEM, the extent of AT enrichment, depletion of GC, G4 level and chromatin accessibility increased from low to high. We also compared OEMs in early and late-replicating genome compartments (Fig.4C), as well as in three different groupings of chromosome size (Fig.4D). In neither analysis was significant differences found in OEM mean or range, suggesting no differential usage of ORIs dependent on timing nor chromosome size. Nonetheless, late replicating areas of the genome were enriched in IZs with low nucleosome occupancy (Fig.4C) and, furthermore, IZs showed increasing AT levels, lower GC levels and increasing chromatin accessibility as chromosome size increased (Fig.4D). These data indicate that despite DNAscent-predicted ORIs being more abundant in late replicating compartments of the *L. major* genome, including across the larger chromosomes, the average efficiency of initiation is similar across replication timing compartments. Whether the genome features we detect at DNAscent-predicted ORIs act to direct DNA replication initiation, or are signatures of initiation occurring more frequently, is unknown.

**Figure 4.**
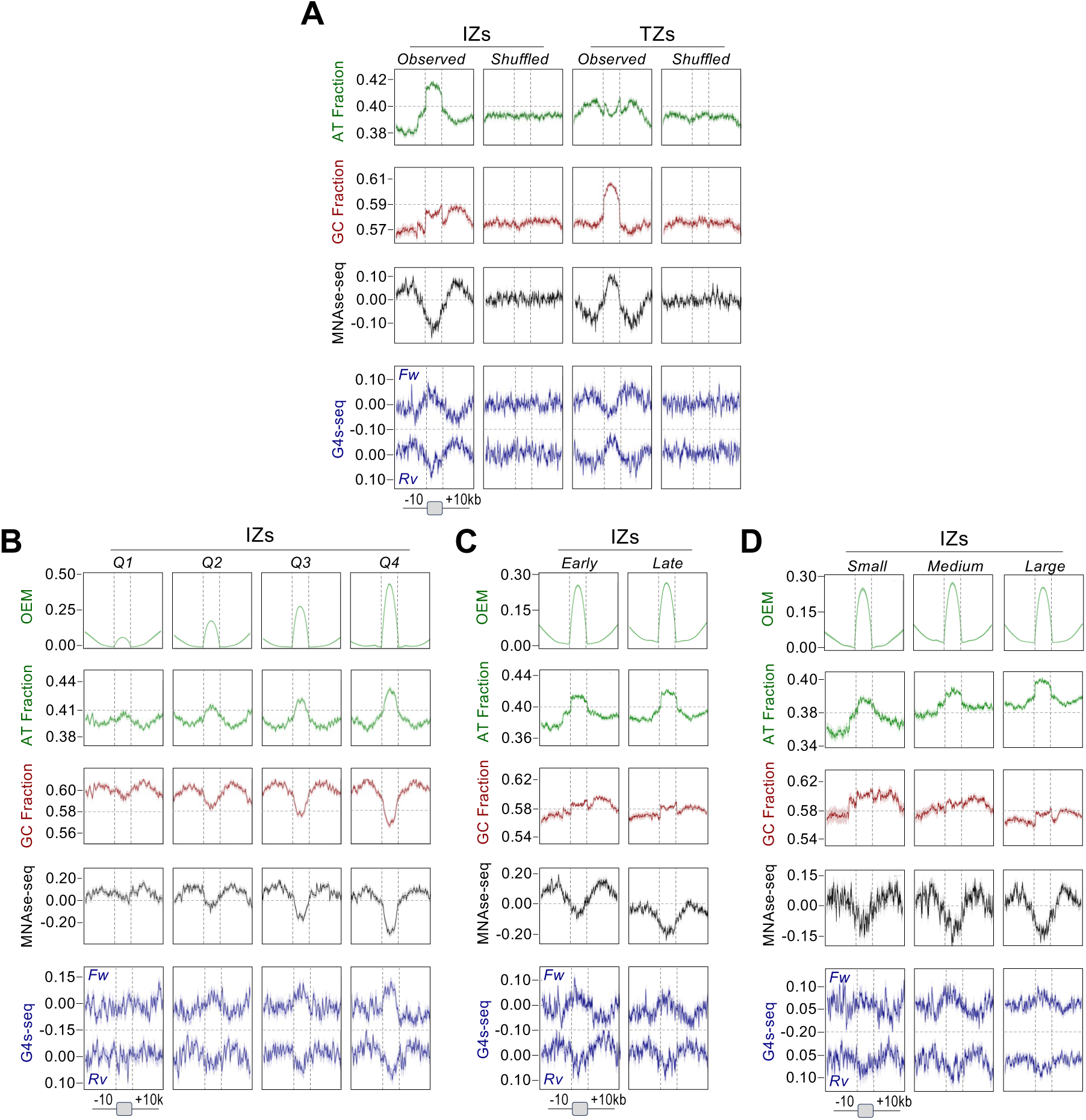
DNA sequence content and chromatin accessibility differs between stochastic initiation and terminations sites and varies according to initiation efficiency. **A)** Metaplots showing global AT and GC content, MNase-seq mapping and G4 profiles around initiation zones (IZs) and termination zones (TZs); the same plots are shown around the same regions after being randomly shuffled (as controls). **B)** Same analysis as in A, but with IZs separated into quartiles (Q1 to Q4) defined according to their average OEM values. **C)** Same analysis as in A, but between IZs found in early and late replicating regions of the genome, as determined by MFA-seq. **D)** Same analysis as in A, but between IZs found in distinct chromosome groups according to their length: small, 0.27 to 0.62 Mb; medium, 0.63 to 0.84 Mb; large, 0.91 to 2.68 Mb.

Next, we asked if the DNA replication initiation sites predicted by DNAscent correlate with origins mapped by SNS-seq^57^ (Fig.S8). Only around 20% of DNAscent ORI midpoints showed an overlap with origins mapped by SNS-seq (Fig S8A). Furthermore, metaplot analysis of the two datasets showed little overlap between density of ORIs detected by DNAscent and SNS-seq origins, with predominant enrichment being more often offset from each other either upstream or downstream (Fig.S8B). Also, we did not detect the strongest enrichment of IZs predicted by DNAscent at sites of transcription termination (convergent or head-tail SSRs, Fig.7B), as seen in SNS-seq mapping^57^, but instead observed a clearer correlation between IZ enrichment and replication timing (Fig. 6, discussed below). Finally, we have recently found that SNS-seq density increases as chromosome size decreases^63^, which is the opposite chromosome length correlation to that we detect here for DNAscent ORIs (Fig.3). Taken together, these analyses suggest that DNAscent predicts a larger and more widely distributed number of initiation events than population-level SNS-seq mapping.

### IZs are regions with reduced levels of nascent RNA, whereas TZs are associated with increased levels of nascent RNA

The near ubiquitous use of multigenic transcription potentially represents a particular problem for genome duplication in kinetoplastids: the need for RNA polymerase (Pol) II to continuously traverse long genomic distances may result in regions of pronounced collisions with the replisome. In both *T. brucei*^32^ and *L. major* (here and ^38,53^), MFA-seq and now DNAscent indicates that constitutive DNA replication initiation, activated early in S-phase is limited to the transcription start or stop sites of polycistronic transcription units (PTUs). To date, no work has asked if and how transcription and replication might intersect within PTUs in *Leishmania*. To address this question, we intersected the RFD profiles derived from DNAscent with annotated transcription direction, and thereby identified genomic segments where DNA replication and transcription travel in the same direction (co-directional), as well as segments where DNA replication and transcription are in opposition (head-on) (Fig. 5A). We did not observe any significant difference between sizes of co-directional and head-on regions (Fig. 5B), indicating a lack of selection for or against one or the other arrangement. Furthermore, we took advantage of recent PRO-seq analysis conducted by Grunebast et al (BioRXiv 10.1101/2023.11.23.568479) and observed that levels of nascent RNA transcripts are similar in co-directional and head-on segments (Fig. 5C). Altogether, these data suggest that the genome organization in this parasite did not evolve to favour co-directional movement of the DNA replisome and transcription machinery and, moreover, DNA replication direction does not seem to influence transcription initiation efficiency. However, visual inspection showed that PRO-seq levels were not uniform across a PTU and there appeared to be a potential correlation between areas of decreased and increased PRO-seq relative to predicted IZs and TZs, respectively (Fig.5D). To test this prediction, we performed metaplot analysis of PRO-seq signal centred on all predicted IZs or TZs and 20 kb of surrounding sequence (Fig.5E).

**Figure 5.**
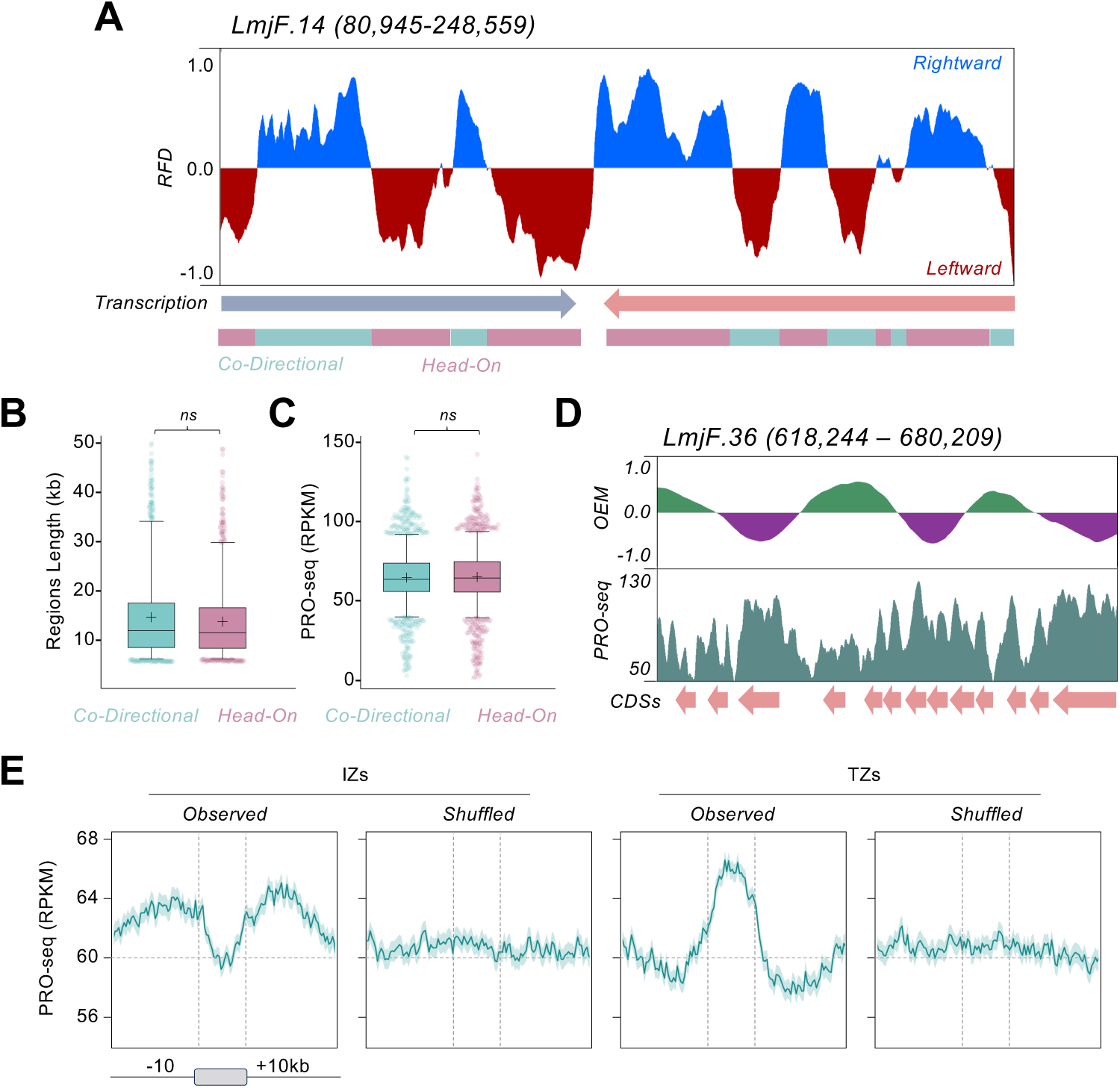
Stochastic DNA replication initiation is associated with lower levels of nascent RNA. **A)** A representative snapshot of *L. major* chromosome 4, showing DNA replication fork direction (from DNAscent; RFD, top) and transcription direction (middle), revealing regions (bottom) where DNA replication and transcription move in the same direction (co-directional) or in opposition (head-on). **B)** Comparing lengths of co-directional and head-on regions across the genome. **C)** Comparing levels of nascent RNA (measured by PRO-seq; Grunebast et al, BioRXiv 10.1101/2023.11.23.568479) in co-directional and head-on regions across the genome. For both C and D, a Kruskal-Wallis test was applied (ns: no significant difference). **D)** Representative snapshot of chromosome 36 comparing origin efficiency metric (OEM) profile and levels of nascent RNA. **E)** Metaplots showing global levels of nascent RNA in initiation zones (IZs) and termination zones (TZs); profiles around the same zones after being randomly shuffled are shown as controls.

This analysis revealed a clear decrease in PRO-seq signal within the IZs and an increase within the TZs. Taken together, these data suggest that transcription and DNA replication do intersect within PTUs, with sites of reduced RNA Pol II transcript level corresponding with stochastic DNA replication initiation, and the inverse correlation between areas of increased transcripts and stochastic DNA replication termination.

### Genome variation correlates with DNA replication timing

To ask if the organisation of *L. major* DNA replication as small numbers of constitutive initiation sites and much larger numbers of stochastic ORIs correlates with replication timing and mutation patterns, we compared the OEM profiles from different genome compartments: the single early-replicating SSR in each chromosome, late-replicating SSRs, subtelomeres, and within and around the CDSs of PTUs (Fig. 6A). The DNA replication timing of each of these loci broadly correlated with initiation efficiency, with early-replicating SSRs having the highest OEMs, and the three other loci displaying lower and similar OEMs (Fig.6B and 6C). At each of these loci we then assessed mutation levels by comparing the number of SNPs that form there after growth in culture. The highest levels of SNP accumulation were seen at early replicating SSRs, where the highest OEM signals were seen (Fig.6D and 6C), indicative of frequent DNA replication initiation and consistent with constitutive activation. At both late-replicating SSRs and subtelomeres, some SNP accumulation was also seen, matching the lower OEMs and the fact that a substantial proportion of these loci appeared to be TZs and not IZs (Fig.6D and 6C). At many CDSs, OEM analysis indicated asymmetric DNA replication activity, with IZs and TZs predicted at one side or the other, an effect that was reflected in very modest but nonetheless detectably asymmetric accumulation and loss of SNPs (Fig.6D and 6C). Taken together, these data indicate that every DNAscent-predicted locus of DNA replication initiation is prone to increased mutagenesis and, moreover, the level of such mutagenesis reflects the measured efficiency of DNA replication initiation.

**Figure 6.**
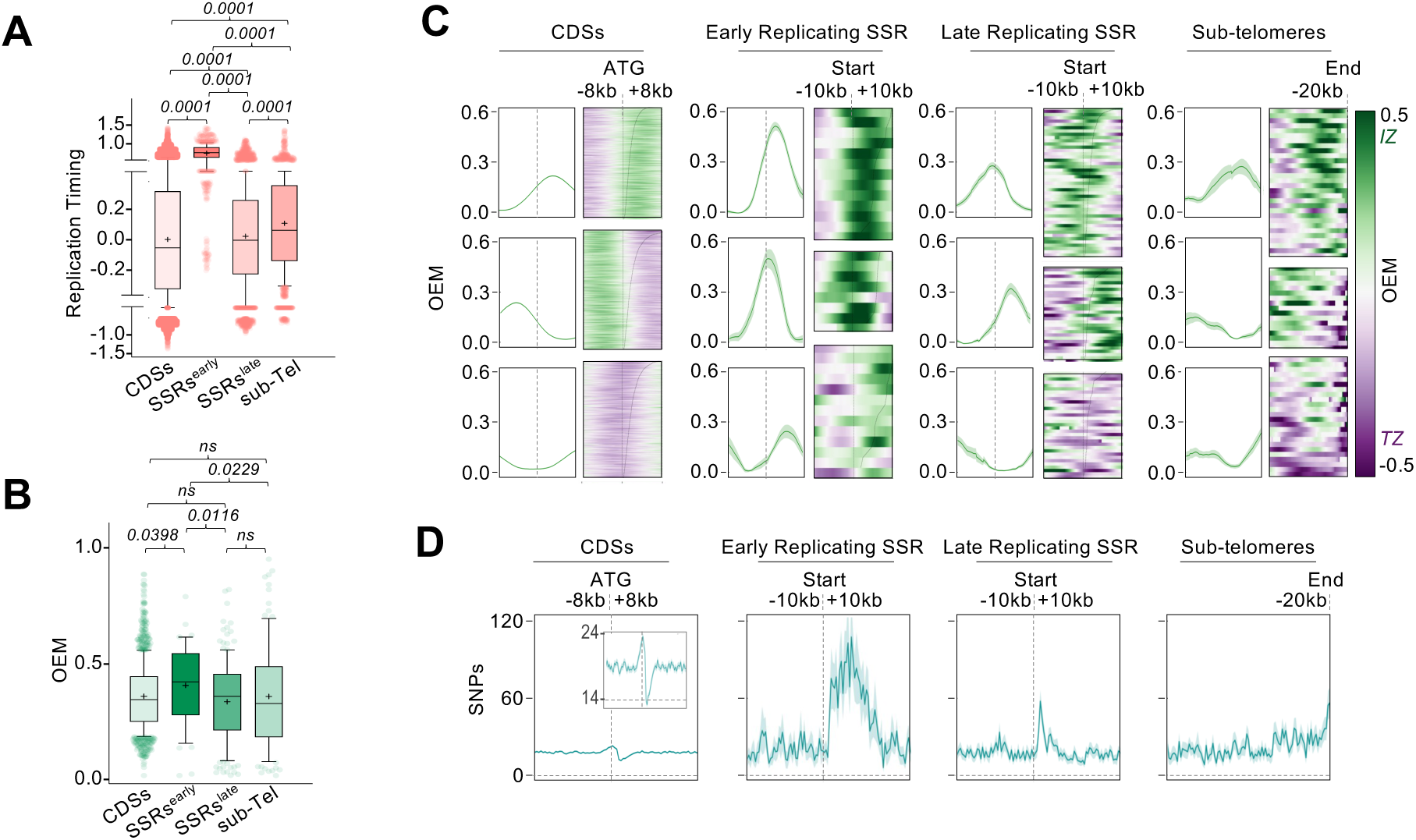
Mutation levels at site of DNA replication initiation correlate with initiation efficiency and timing. **A)** DNA replication timing and **B)** positive origin efficiency metrics (OEMs) are shown for four genomic regions: coding sequences (CDSs), early-replicating SSRs, late-replicating SSRs, and chromosome subtelomeres (sub-Tel); p values from Kruskal-Wallis tests are indicated (ns: not significant). **C)** Metaplots of global OEM values and **D)** SNP density around the same genomic regions.

## Discussion

Analysis of nuclear DNA replication in the ubiquitous human and animal pathogen *Leishmania* has, to date, relied on population-level next generation sequencing approaches ^38,53,57^. Here, we have made use of the capacity of Nanopore sequencing to detect the nucleotide analogue BrdU in long sequence reads, first described by Muller and colleagues^8^, to perform single molecule analysis of DNA replication in *L. major* and understand how this eukaryotic parasite copies its genome. Our work not only confirms the unprecedented use of just a single constitutive locus of DNA replication initiation in each *L. major* chromosome, but reveals thousands of previously undetected, highly stochastic initiation events that escaped detection by all previous approaches. Furthermore, we show that the use of abundant, stochastic replication initiation events relates to the unusual chromosome size-dependent timing of *L. major* DNA replication. Finally, we show that all predicted sites of DNA replication initiation are marked by localised patterns of base content, chromatin organisation, transcription and mutation, revealing that that the unusual, bimodal programme for DNA replication adopted by *Leishmania* has shaped the evolution of the parasite’s genome.

The first attempt to map origins in *Leishmania* species relied on MFA-seq, an approach that in *S. cerevisiae* and relatives shows considerable consistency with locations of conserved origins^3,68–70^, and in *T. brucei* clearly correlates with binding locations of one subunit of ORC^32,33^. In both cases, and in accordance with eukaryotic DNA replication canon, multiple origins were detected per chromosome. In *Leishmania*, however, MFA-seq analysis of two species, *L. major* and *L. mexicana*, was only able to detect a single putative origin of replication in each chromosome^53^. This unprecedented observation was met with caution, primarily because one origin per chromosome, although in theory enough to replicate many of the smaller chromosomes during S-phase, is undoubtedly insufficient to completely replicate the larger parasite chromosomes^27,31,53^. Indeed, we hypothesised that other origins must be present but are not used constitutively (and thus eluded MFA-seq detection), or the parasite relies on other mechanisms, such as DNA repair, to direct DNA replication at other sites in the genome. Indeed, further refinement of our MFA-seq approach suggested that subtelomeric replication may occur outside S-phase and is dependent on components of the 9-1-1 DNA checkpoint clamp (involved in DNA repair)^38^. Studies using other methodologies have challenged the MFA-seq findings^30,55^, with the only other genome-wide analysis, SNS-seq, predicting ∼5000-6000 putative origins and limited correspondence to MFA-seq data^57^. Thus, complete understanding of how *L. major* replicates its genome remained elusive. DNAscent^8,61^ is a methodology that detects DNA replication in single DNA molecules and allows the prediction of fork direction, leading to predictions of initiation and termination loci across the genome. DNAscent has a number of advantages over previous approaches applied to *Leishmania*: it is much more sensitive than MFA-seq, in that it examines long single DNA molecules; it provides for genome-wide analysis, which DNA combing approaches used so far in *Leishmania* do not; and it relies only on the detection of BrdU incorporated into sequenced DNA, thus circumventing the need for processing and enrichment to detect replicating DNA molecules.

We suggest that the use of DNAscent more accurately predicts sites of DNA replication initiation in *L. major* than any previous analysis, based on two broad observations. The first observation is that DNAscent confirms and extends the findings of population-level MFA-seq mapping. Most prominently, BrdU signal density in Nanopore reads, allied to DNAscent analysis of the pattern of BrdU accumulation across the genome, shows that the only loci where signal is uniformly seen in unsynchronised *L. major* promastigote cells correspond to a single SSR in each chromosome, overlapping with MFA-seq mapping (Fig.1, Fig.S1). In addition, HU-mediated G1/S synchronization of the cells followed by release and synchronous progression across S-phase is consistent with coordinated bidirectional replication fork movement from only these loci and with remarkable consistency of movement between chromosomes (Figs.1,2). Finally, RFD predictions suggest that DNA replication initiates from relatively discrete locations within most of these SSRs (Fig.3, Fig.S6). These findings show that DNAscent recapitulates and improves upon MFA-seq data, validating its effectiveness, and indicate that one SSR in each chromosome is the sole site of a coordinated DNA replication initiation in promastigote cells at the onset of S-phase. No study has yet described ORC genomic localisation in the *L. major genome*, and so we cannot say if these early-replicating SSRs are true origins or are instead loci at which wider initiation sites (see below) are concentrated. Nonetheless, a range of data lead us to hypothesise that these early-replicating SSRs are likely to be ORC-defined origins. As we have noted before^53^, a significant fraction of the early-replicating SSRs in *L. major* are syntenic with ORC-bound origins in *T. brucei*. Here, we show that RFD shifts within most SSRs are notably abrupt, and thus appear more comparable to the RFD shifts seen at sequence-defined origins in *S. cerevisiae* ^9,66,67^ than the more diffuse RFD shifts seen in human cells (either because of greater heterogeneity of origin usage or clusters of origins)^66,71^. One study has suggested these SSRs are also the locations of centromeres^54^, which are also found in single copy in each chromosome of *T. brucei* and are notably early replicating^32^, even when they reside in the transcriptionally-silent subtelomeres^34^. Thus, the earliest DNA replication activation event in S-phase of these two related parasites appears to rely on so-far unexplored links with centromere function, which is consistent with BioID analysis indicating proximity of the kinetochore and ORC in *L. mexicana*^72^. Where the parasites appear to differ is in how they programme DNA replication across the genome and cell cycle beyond such centromere-focused origin activity: in *T. brucei*, every SSR binds ORC and around 25% act as origins that can be detected by MFA-seq^32^; in contrast, none of MFA-seq^38,53^, SNS-seq^57^ or DNAscent (this study) provide evidence that SSRs are equivalent, ubiquitous sites of DNA replication initiation in *L. major*.

The second reason for suggesting DNAscent provides a fuller understanding of DNA replication in *L. major* lies in the prediction of more widespread sites of initiation that MFA-seq or any other approach used to date. Theoretical predictions suggest that the relatively small number of ORC-localised origins detected at SSRs by MFA-seq in each *T. brucei* chromosome is around the minimum needed to complete copying of each chromosome in S-phase, and does not provide an excess of potential back-up origins, as seen in yeast, for example^73^. Here, DNAscent indicates that *Leishmania* uses a different strategy to complete replication of each of its chromosomes: the constitutive activation of a single putative origin early in S-phase is supplemented with the abundant use of stochastic ORIs, which are distributed across chromosomes and do not specifically localise to SSRs. The total number of such stochastic *L. major* ORIs is hard to determine from the available data, as such measures will depend on sequence depth. Nonetheless, as DNAscent examines DNA replication patterns on single molecules, it is somewhat comparable to single molecule combing, which previously predicted inter-origin distances in *L. major* of ∼70 kb^57^ and ∼195 kb^55^; here, DNAscent predicts an inter-origin distance of ∼20 kb, which is likely to be close to the minimum detectable by fibre analysis^57^, and suggests the use of substantially larger numbers of ORIs than any previous study has suggested^38,53,55,57^. Most likely, the combination of the sheer number of such ORIs, allied to flexibility in their localisation in the genome, precludes their detection by MFA-seq. In addition, we also find limited overlap between DNAscent and SNS-seq predictions of ORIs in *L. major*, which contrasts with the good correspondence between SNS-seq and Nanopore-BrdU mapping in *P. falciparum* cells undergoing schizogony^22^. Thus, the abundance of stochastic ORIs in *Leishmania* may be truly unusual amongst single-celled eukaryotes. In *S. cerevisiae*, only around 10-20% of origins mapped by BrdU-Nanopore sequencing cannot be aligned with known origins^8,9^, whereas 80% of DNAscent-predicted initiation events in human cells do not match a range of origin prediction approaches^74^. Thus, it is possible that *Leishmania*’s abundant use of stochastic DNA replication initiation may have parallels with at least some metazoans^10^.

From the available data on DNA replication, we can only speculate on the nature of the abundant, non-constitutive ORIs in *L. major*. It is possible that the stochastic ORIs are also designated by ORC binding, and therefore *L. major* has a previously unanticipated abundance of ‘true’ origins. However, ORC designation of such widely dispersed origins may be problematic for *Leishmania*, since in other eukaryotes ORC associates with DNA and recruits the MCM helicase in G1. As most of the DNAscent-predicted ORIs are spread throughout the PTUs of *L. major*, they might be predicted to present a much greater impediment to transcription than in other eukaryotes, given the ubiquitous use of polycistronic transcription. It is possible that ORC might be loaded, perhaps at the single early-replicating SSR, in G1 but is loosely bound and mobile, and so moves throughout the genome until activation of DNA replication in S-phase, which could then occur highly flexibly, wherever ORC is found. Alternatively, and has been suggested by Lombraña and colleagues^57^, ORC recruitment of MCM might lead to a stable pre-RC complex only at the constitutive origins, with MCM in other locations free to move away from ORC and lead to initiation. Both above suggestions are consistent with the lack of any conserved sequence features of the stochastic ORIs, and the shifts in localised base and chromatin content, as well as native transcript levels, we observe might reflect loci where the replication machinery lingers in the genome. Finally, is it possible that stochastic initiation events are ORC-independent. Such activator-independent initiation of DNA replication has been described eukaryotes but is normally only readily detected as a ‘back-up’ reaction after mutation of the activator-origin machinery, or *in vitro*^75–79^. How, then, *Leishmania* might programme the use of both ORC-dependent and -independent reactions into the normal course of DNA replication is unclear. We have previously reported subtelomeric DNA replication in *L. major* that appears to not be limited to S-phase and is dependent on Rad9^38^, but whether this might also account for all stochastic DNA replication reactions is unclear.

Whatever the nature of the abundant, stochastic ORIs detected by DNAscent, a further question raised by this study is why *Leishmania* has evolved a DNA replication programme that combines use of a single, constitutive site of replication initiation per chromosome supplemented with much more abundant, stochastic ORIs. An explanation may lie in directing patterns of genome change. Here we show that SNPs accumulate at all sites of DNA replication initiation, but these are most pronounced at the single constitutive origin in each chromosome, confirming and extending our previous analyses^38,39^. The same effect has been observed at human origins, with patterns of mutation differing in ‘core’ and cell-type specific origins^80^. *Leishmania* may then have evolved to spread such initiation-induced mutagenesis across the genome, rather than focusing it at a small number of sites, as this could promote adaptation.

Intriguingly, recent work using MFA-seq mapping has revealed distinct levels of predicted origins in the highly transcribed core and largely untranscribed subtelomere compartments of the *T. brucei* genome, with the latter compartment having MFA-seq peaks and displaying notably greater instability^34^. No work has applied DNAscent in *T. brucei*, and so it remains unclear if subtelomere DNA replication may rely on similar stochastic origins that are found genome-wide *in L. major*. Nonetheless, the data we present here provides a mechanistic link between DNA replication programming and genome plasticity in *Leishmania*, which may have parallels with other kinetoplastids.

## Methods

### Parasite culturing and BrdU labelling

Promastigotes derived from *Leishmania major* V9 (MHOM/IL/80/Friedlin) strain were cultured at 26 °C in HOMEM medium supplemented with 10% heat-inactivated foetal bovine serum. Prior to labelling, parasites were seeded at 5x10^5^ cells.mL^-1^ and allowed to proliferate until exponentially growing at 5x10^6^ cells.mL^-1^. BrdU was added to the culture to a final concentration of 150 µM. After 2 minutes of incubation at 26 °C, cells were collected by centrifugation at 2,300 g for 2 minutes. Cell pellets were immediately resuspended in culturing medium containing 1 mM thymidine and incubated at 26 °C for 1 hour. Then, cells were collected by centrifugation at 2,300 g for 2 minutes and pellets were stored at −20 °C until used. The same labelling approach was used for synchronised cells after treatment with hydroxyurea (as described in the main text).

### High molecular weight DNA extraction

To ensure isolation of long DNA molecules, genomic DNA extractions were performed using the MagAttract High Molecular Weight DNA Kit (QIAGEN). For each extraction, a pellet containing approximately 5x10^8^ cells was removed from −20 °C storage and immediately resuspended in lysis buffer. All further processing steps were performed following the manufacturers’ instructions. After elution, DNA samples were incubated at 4 °C for 24 to 48 hours to allow complete sample homogenisation.

### Oxford Nanopore Technology GridION sequencing

High molecular weight genomic DNA samples were subjected to library preparation for Nanopore sequencing using the Ligation Sequencing Kit SQK-LSK110 (Oxford Nanopore Technologies). Approximately 5 μg input genomic DNA was used in each reaction and all processing steps were performed following the manufacturers’ instructions. Libraries were loaded onto R9.4.1 GridION flow cells (Oxford Nanopore Technologies) and sequenced for up to 48 hours. When needed, sequencing was paused, flow cells washed and reloaded with extra library material.

### Processing of Nanopore sequencing files

The Nanopore run directories were processed with *Guppy basecaller v6.4.2* with configuration *dna_r9.4.1_450bps_fast.cfg*. The resulting fastq files were aligned to the reference genome *Leishmania major Friedlin* v45 (https://tritrypdb.org/tritrypdb/app) using *minimap2* ^81^ with setting *-x map-ont*. The resulting bam files were sorted and indexed with *samtools* ^82^. Sorted bam files were subjected to BrdU calling with the *DNAscent detect* function from *DNAscent v2* ^61^ (https://github.com/MBoemo/DNAscent-commit7e4be09) using commands *index, detect* and *forkSense*. Output files from *DNAscent detect* contained the probability of BrdU at each thymidine in the genome.

These files were rearranged to a bedgraph-like format containing six columns: chrom, start, end, percentage of reads with BrdU calls, read depth and number of reads with BrdU calls. The output from *DNAscent forkSense* contained coordinates of left- and right-moving forks as well as for the ORI and TER sites. These files were also rearranged to a bedgraph-like format containing nine columns: chrom, start, end, probability of left-moving fork, probability of right-moving fork, read name, strand, alignment start and alignment end. This workflow was run within a mamba environment and implemented as a Snakemake ^83^ pipeline as detailed here: https://github.com/glaParaBio/dnascent-fork-detection.

### Generation of BrdU scores

Output files from *DNAscent detect* were processed to remove all BrdU calls with probability <0.5 whilst retaining BrdU calls with probability ≥0.5. Retained BrdU calls were averaged into 500 bp sequential windows across the genome using *bedtools MapBed* on *Galaxy* (https://usegalaxy.org/)^84^. The scores from the resulting bedgraph files were converted into z-scores calculated in rolling windows of 15 kb by using R.

### Generation of ORI densities files

Given the BrdU pulse length (2 min) and the average fork speed in *L. major* (2.5 kb.min^-1^), we expect ORIs fired immediately before labelling to have have maximum length of 10 Kb. Therefore, output files from *DNAscent forkSense* were processed to retain only ORI calls ≤ 10 kb. Midpoints for retained ORI calls were determined and were averaged into 500 bp sequential windows across the genome using *bedtools MapBed* on *Galaxy.* These are the *ORI density* files. The *ORI density* bedgraph files were processed by calculating z-scores calculated in rolling windows of 30 kb using R. These are the *Relative ORI density* files.

### Generation of replication forks files

Output files from *DNAscent forkSense* were processed to remove all fork calls with probability < 0.5 whilst retaining fork calls with probability ≥ 0.5 in reads with length of at least 15 kb. Retained forks calls were averaged into 20 bp sequential windows across the genome using *bedtools MapBed* on *Galaxy*. In this way, two bedgraph files, one with the right-moving forks and another with the left-moving forks, were generated.

### Replication Fork Directionality (RFD) profiles

Bedgraph files with the right- and left-moving forks were used as input for determining RFD in 1kb rolling windows. RFD has been previously defined ^65,66,71^ and was calculated as ((right-moving forks – left-moving-forks)/(right-moving forks + left-moving-forks)).

### Origin Efficiency Metrics (OEM) profiles

Bedgraph files with the right- and left-moving forks were used as input. OEM was calculated in 10 kb rolling windows divided into 5 kb left-windows and 5 kb right-windows as previously defined ^65–67^. Calculations were performed as the following: [(right-moving forks)/(right-moving forks + left-moving-forks)]*^Left-window^* - [(right-moving forks)/(right-moving forks + left-moving-forks)]*^Right-window^*_._

### SNP density

Sequencing files of *Leishmania major* strain LT252 (MHOM/IR/1983/IR) were obtained from ^39^. SNPs were called using *FreeBayes* on *Galaxy*. Only SNPs with at least two supporting reads and with QUAL > 30 were retained and averaged into 1 kb sequential windows across the genome using *bedtools MapBed*.

### Heatmaps, metaplots and graphs

Heatmaps and metaplots were generated with *deepTools plotHeatmap* and *plotProfile* tools, respectively, on Galaxy. Remaining graphs and their respective statics analysis were generated using Prism GraphPad.

## Data availability

ONT sequences used for DNAscent analysis are available at the Eurpoean Nucleotide Archive under accession number PRJEB82099.

## Acknowledgments

We thank Mike Boemo for helpful discussions about establishing DNAscent in *Leishmania*, and all current and previous members of the McCulloch lab for input. This work was supported by the Wellcome Trust [224501/Z/21/Z], the BBSRC [BB/N016165/1, BB/R017166/1, BB/W001101/1], the MRC [MR/S019472/1], and the European Union’s Horizon 2020 research and innovation programme under the Marie Sklodowska-Curie grant agreement No 750259 [Individual Fellowship, RECREPEMLE]. The Wellcome Centre for Integrative Parasitology was supported by core funding from the Wellcome Trust [104111]. Parts of Figure 1A were generated using BioRender.

---

**Figure S1.**
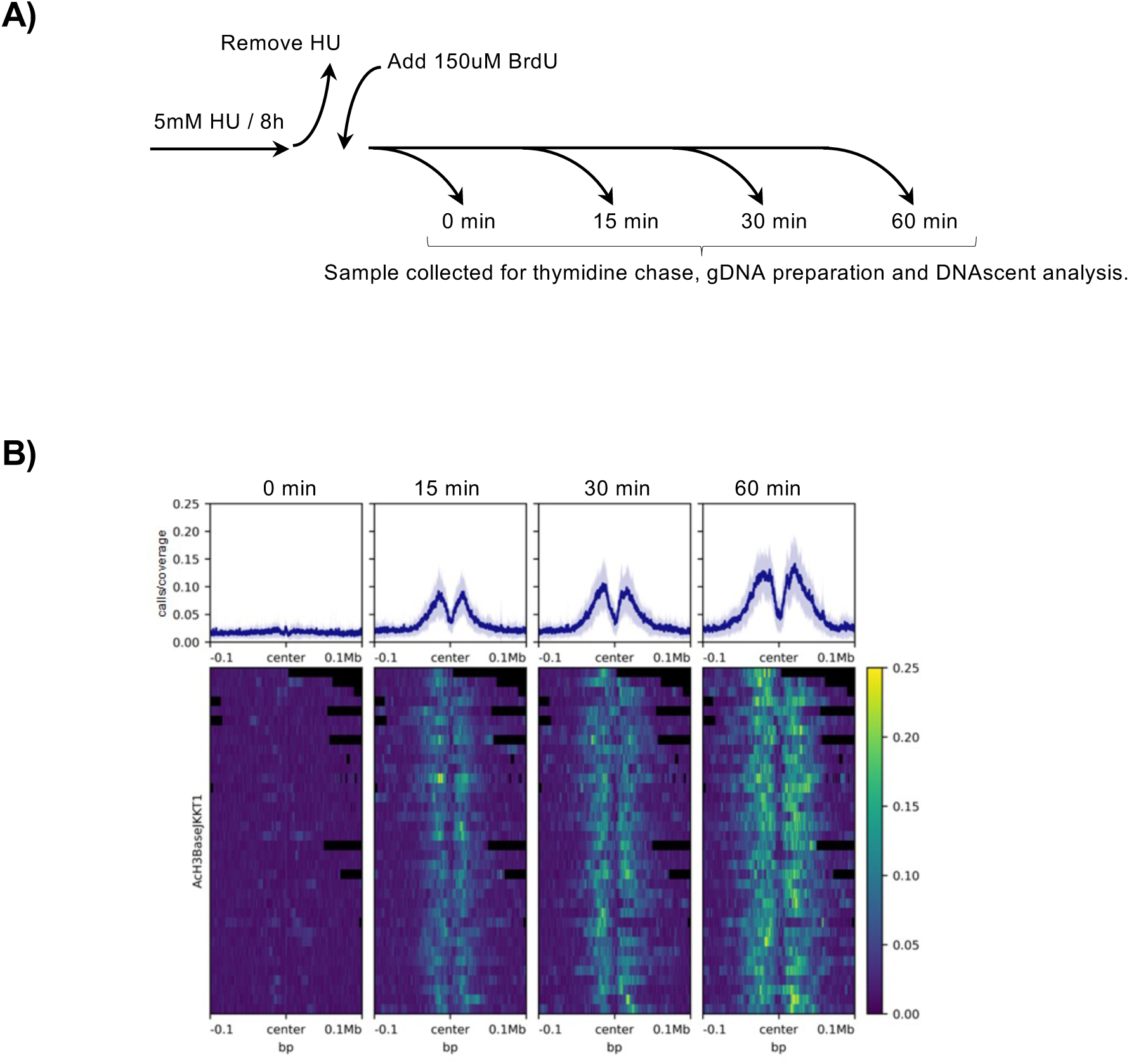
Testing incorporation of BrdU into *Leishmania major* promastigote nuclear DNA. **A)** Schematic of the experimental approach. Exponentially growing *L. major* promastigotes were treated with HU for 8 hrs, then moved into fresh HU-free medium containing 150 uM BrdU and cells sampled 0, 15, 30 and 60 min later. For each sample, 1 mM thymidine was added for 1 hour, then high molecular weight DNA was extracted and subjected to Oxford Nanopore Technologies sequencing. **B)** DNAscentv2 analysis of BrdU levels in the above sequences, with data shown as metaplots of signal around all early-replicating SSRs (predicted by MFA-seq^53^).

**Figure S2.**
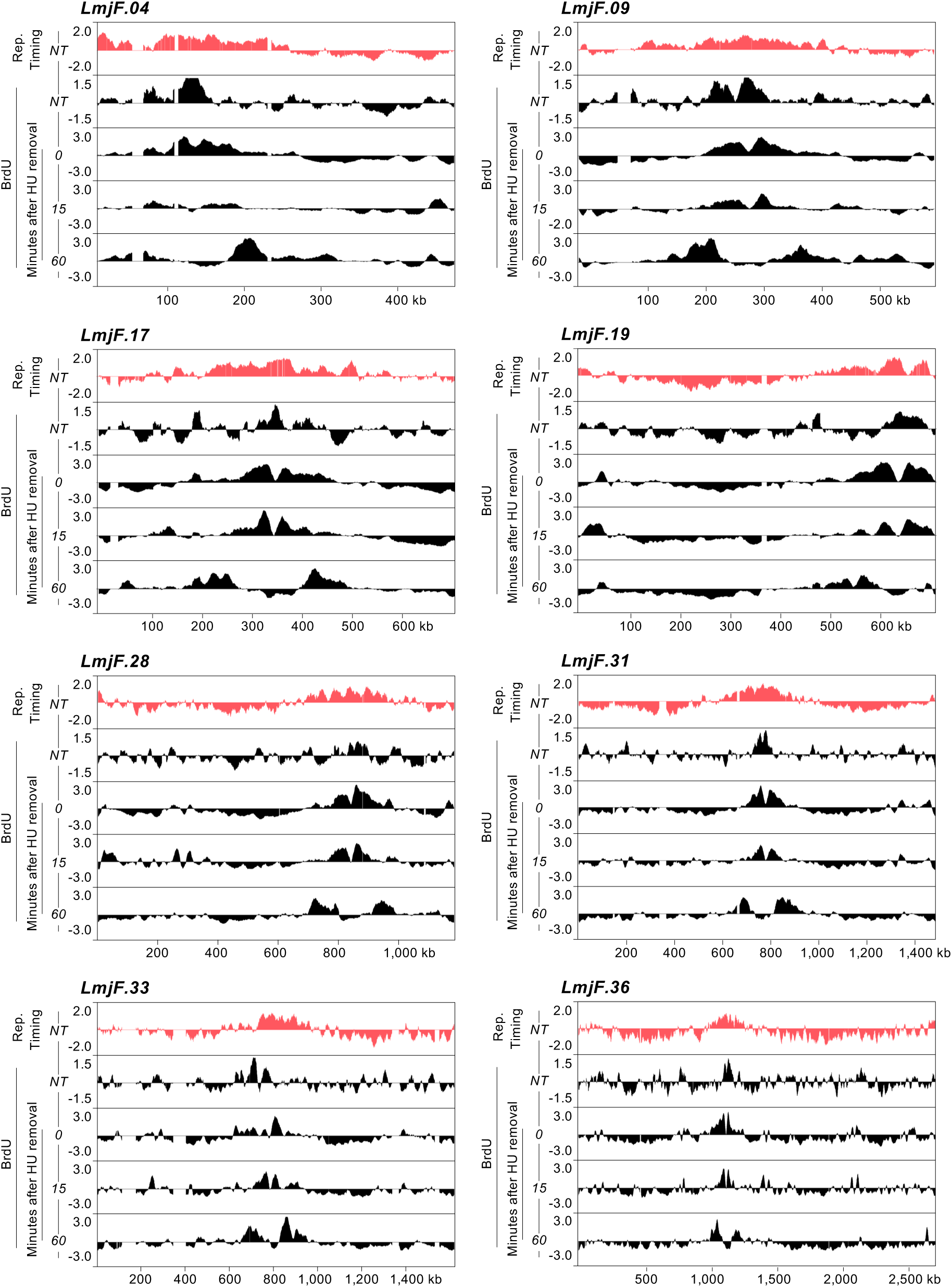
Mapping BrdU incorporation profiles with DNAscent confirms centromeric SSR-driven DNA replication initiation. BrdU scores (black) across eight *L. major* chromosomes, comparing unsynchronised cells (NT) and cells 0, 15 and 60 minutes after release from HU cell cycle stall. The top track (salmon) shows the DNA replication profile in the same chromosomes as determined by MFA-seq.

**Figure S3.**
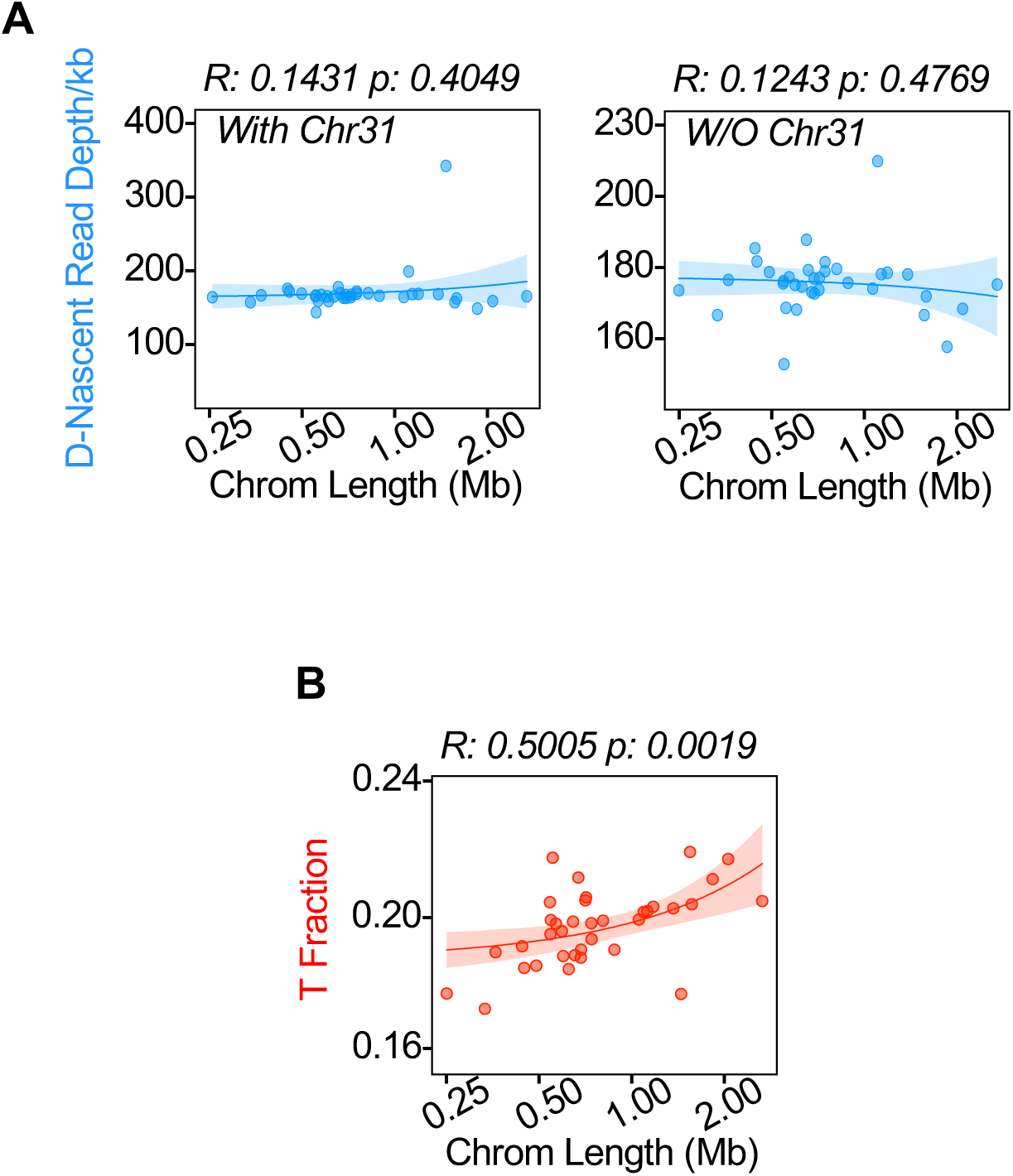
DNAscent output is not influenced by Nanopore read depth or by base T content of *L. major* chromosomes. **A)** Regression analysis comparing average read depth in each chromosome relative to its length, either including (left) or excluding (right) chromosome 31, which is estimated to be tetraploid. B) Regression analysis comparing the fraction of bases that are T in each chromosome relative to its length.

**Figure S4.**
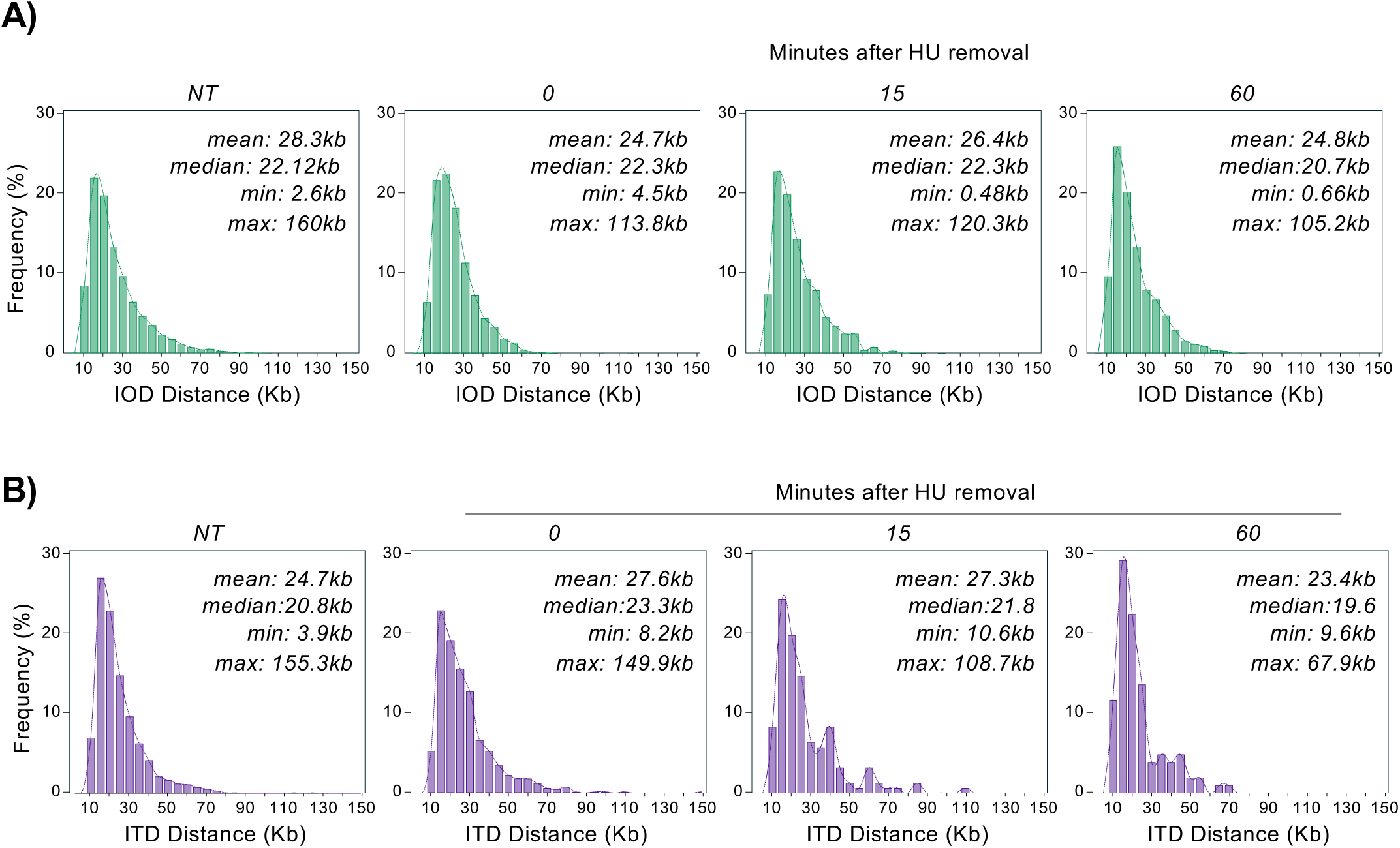
DNAscent reveals the density of predicted initiation and termination sites in the *L. major* nuclear genome. Plots show the range of distances between every DNAscent-predicted initiation site (inter ‘origin’ distance, IOD) and between every predicted termination site (inter-termination distance, ITD) in unsynchronised cells (NT) and cells 0, 15 and 60 minutes after release from HU cell cycle stall.

**Figure S5.**
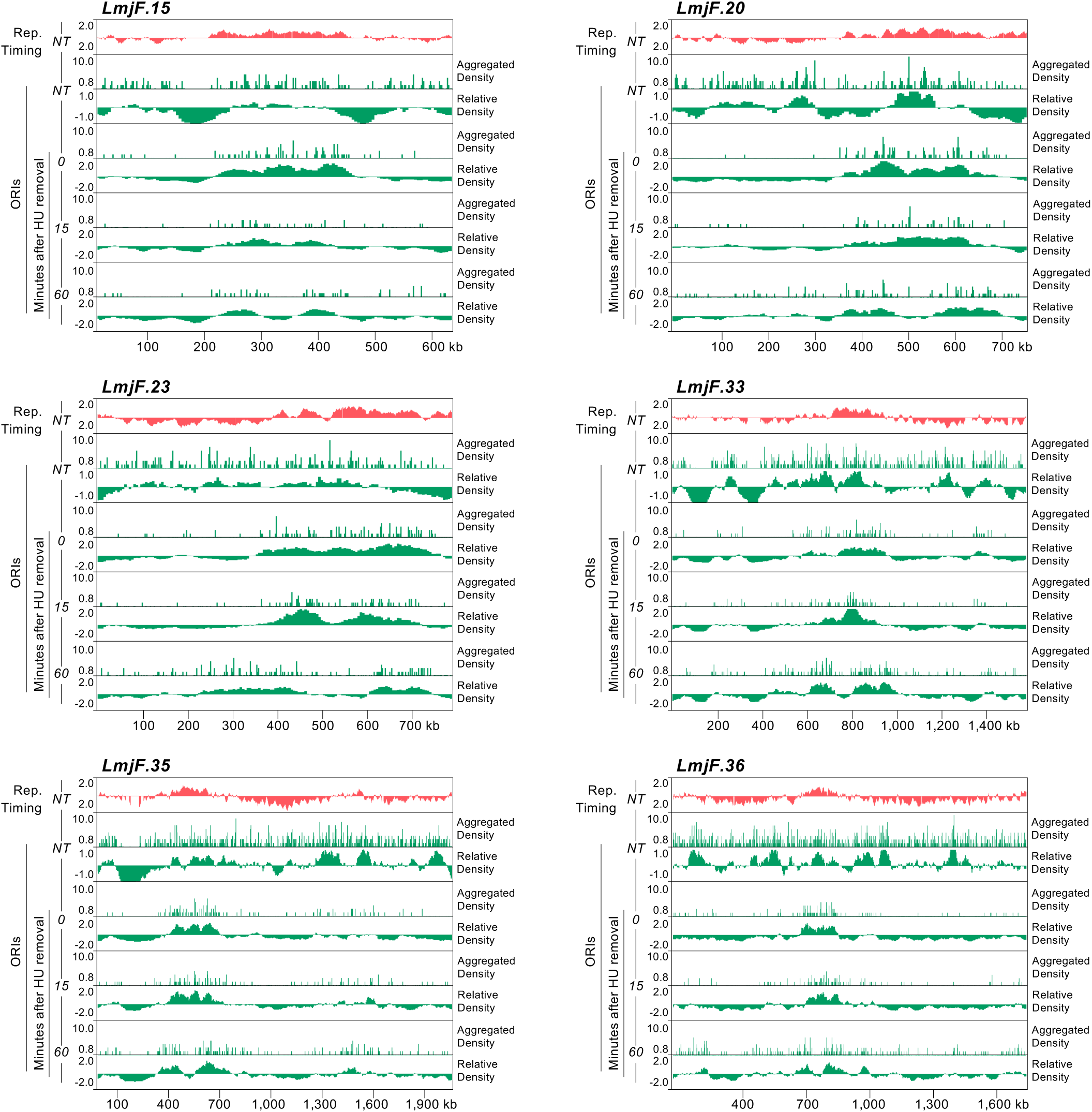
DNAscent shows that a single centromeric SSR in each chromosome is activated in early S phase and reveals further, widespread initiation events. **A)** Snapshot showing DNAscent-predicted ORI density (green, aggregated and relative) across six *L. major* chromosomes in unsynchronised (NT) cells and in cells 0, 15 and 60 mins after release from HU stall; the top track (salmon) shows the DNA replication profile as determined by MFA-seq.

**Figure S6.**
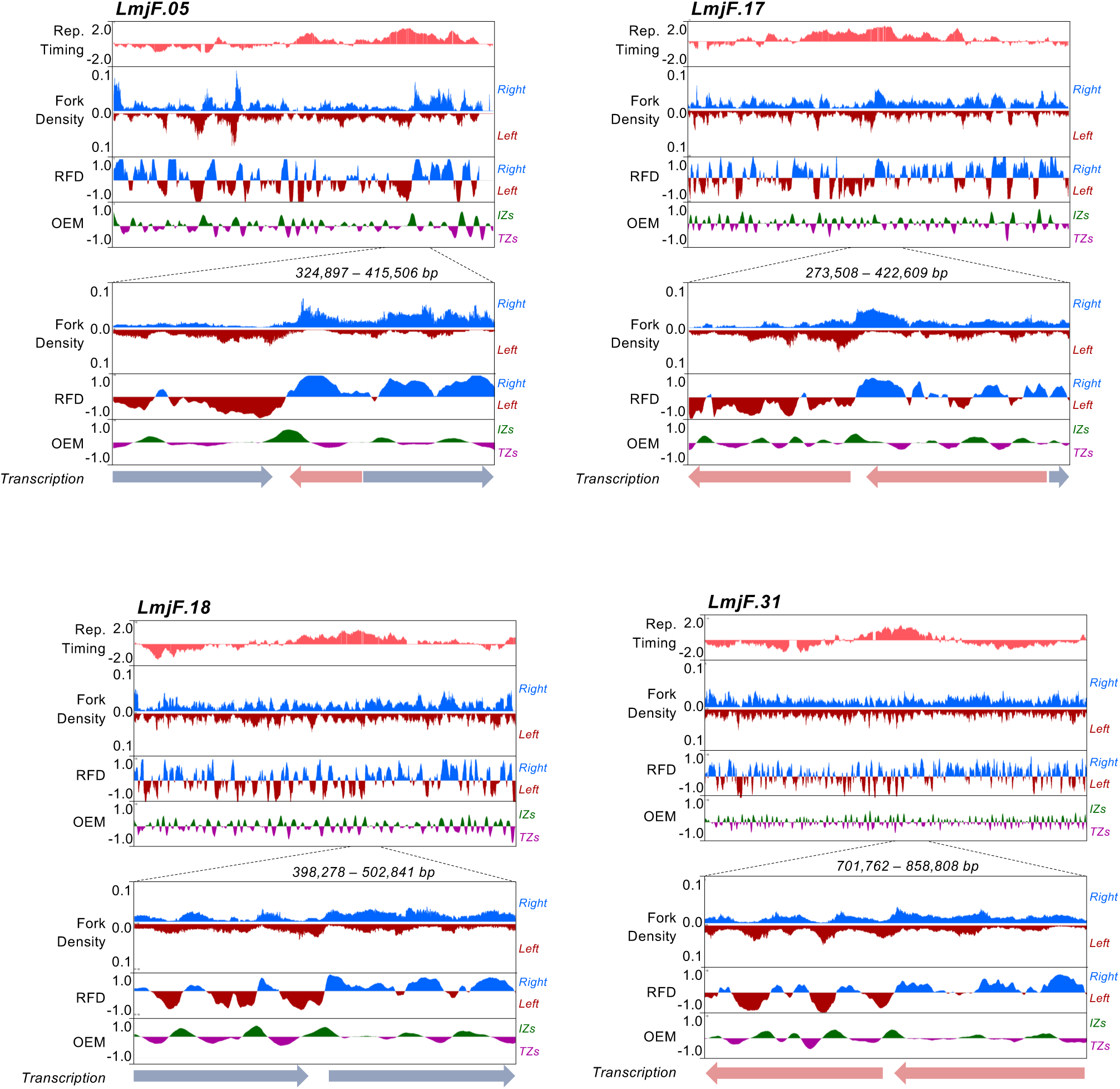
Analysis of DNA replication forks detected by DNAscent. DNA replication fork density and quantification of predicted replication fork direction (RFD) and origin efficiency metrics (OEM) is shown for four chromosomes, in each case across the whole chromosome and (as a zoom) in the regions surrounding early-replicating SSRs (predicted by MFA-seq; organisation of PTUs around the early-replicating SSRs is shown below the plots).

**Figure S7.**
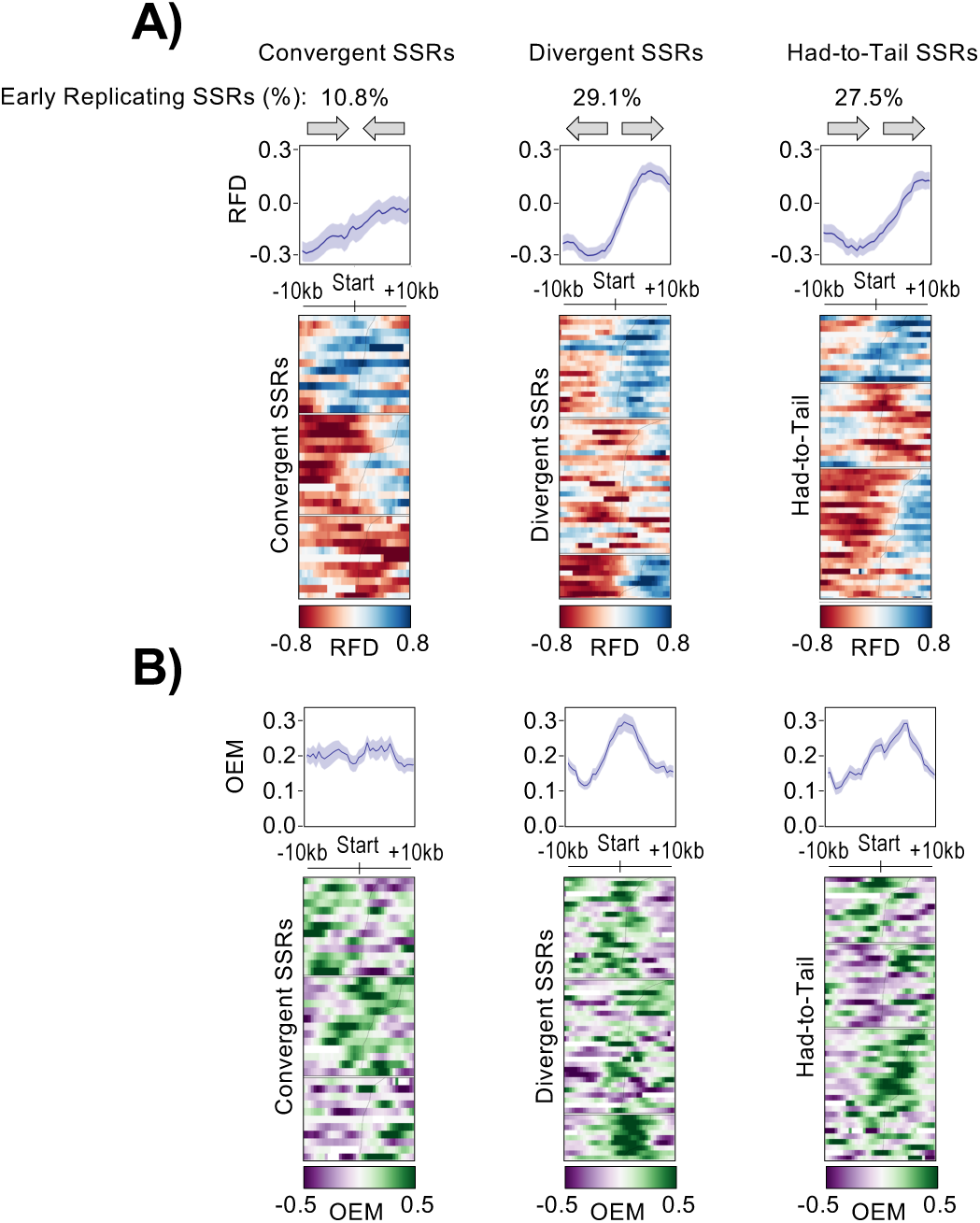
DNAscent analysis of DNA replication around all SSRs. **A)** SSRs were separated into those where surrounding polycistronic transcription terminates (convergent SSRs), where bidirectional transcription initiates (divergent SSRs), or where upstream transcription terminates and downstream transcription initiates (Head-to-Tail SSRs). DNAscent-predicted replication fork direction (RFD) is shown around each SSR (lower diagram), or as a metaplot of all SSRs (upper panel). The fraction of SSRs that are predicted by MFA-seq to be early-replicating in each category is shown. **B)** As in (A) but showing origin efficiency metrics (OEMs).

**Figure S8.**
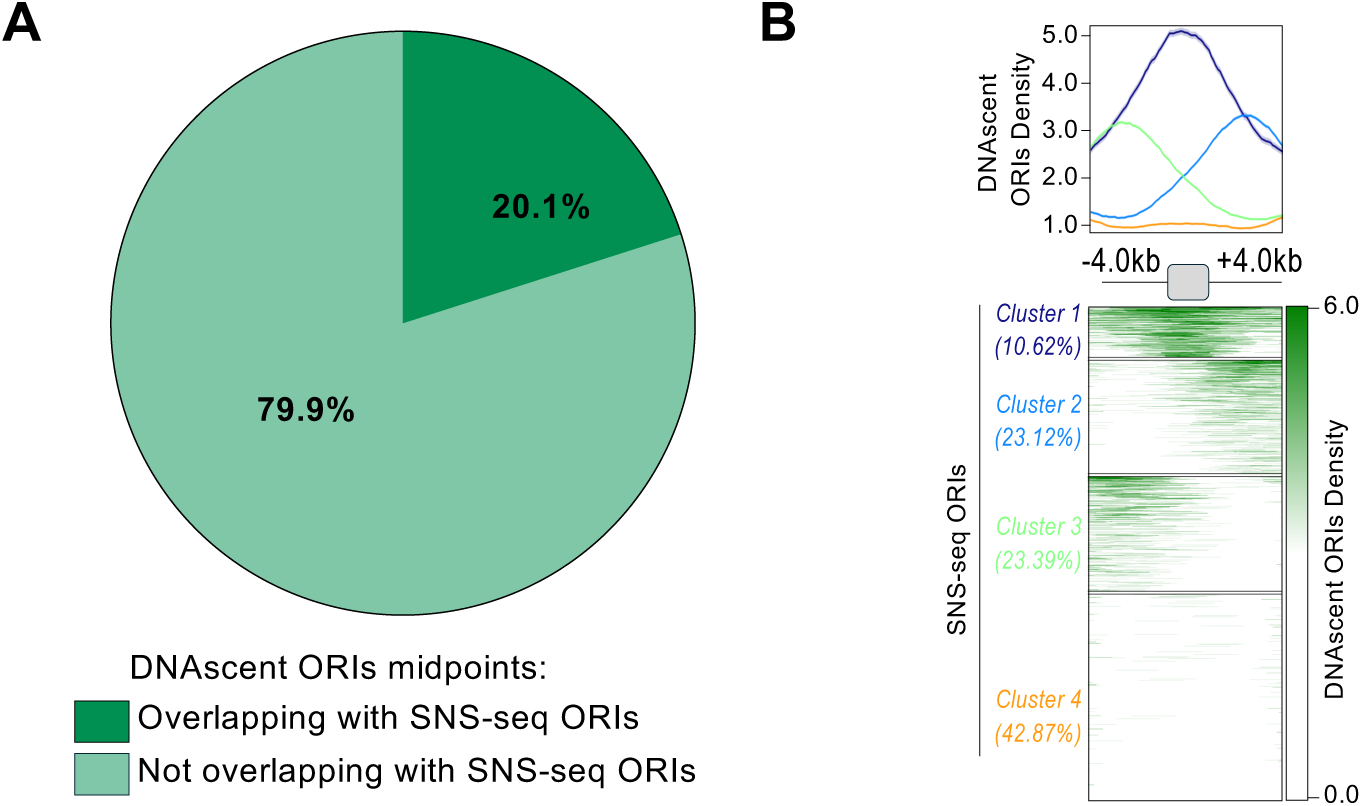
Little correlation between DNAscent prediction of DNA replication and SNS-seq. **A)** Pie-chart showing the proportion of DNAscent predicted ORIs where the midpoint overlaps with SNS-seq predicted ORIs. **B)** Metaplots showing density of DNAscent-predicted ORIs around all SNS-seq signal peaks.

