## Supplementary figures and images for "Nuclear DNA replication in *Leishmania major* relies on a single constitutive origin per chromosome supplemented by thousands of stochastic initiation events"

### Supplementary figures

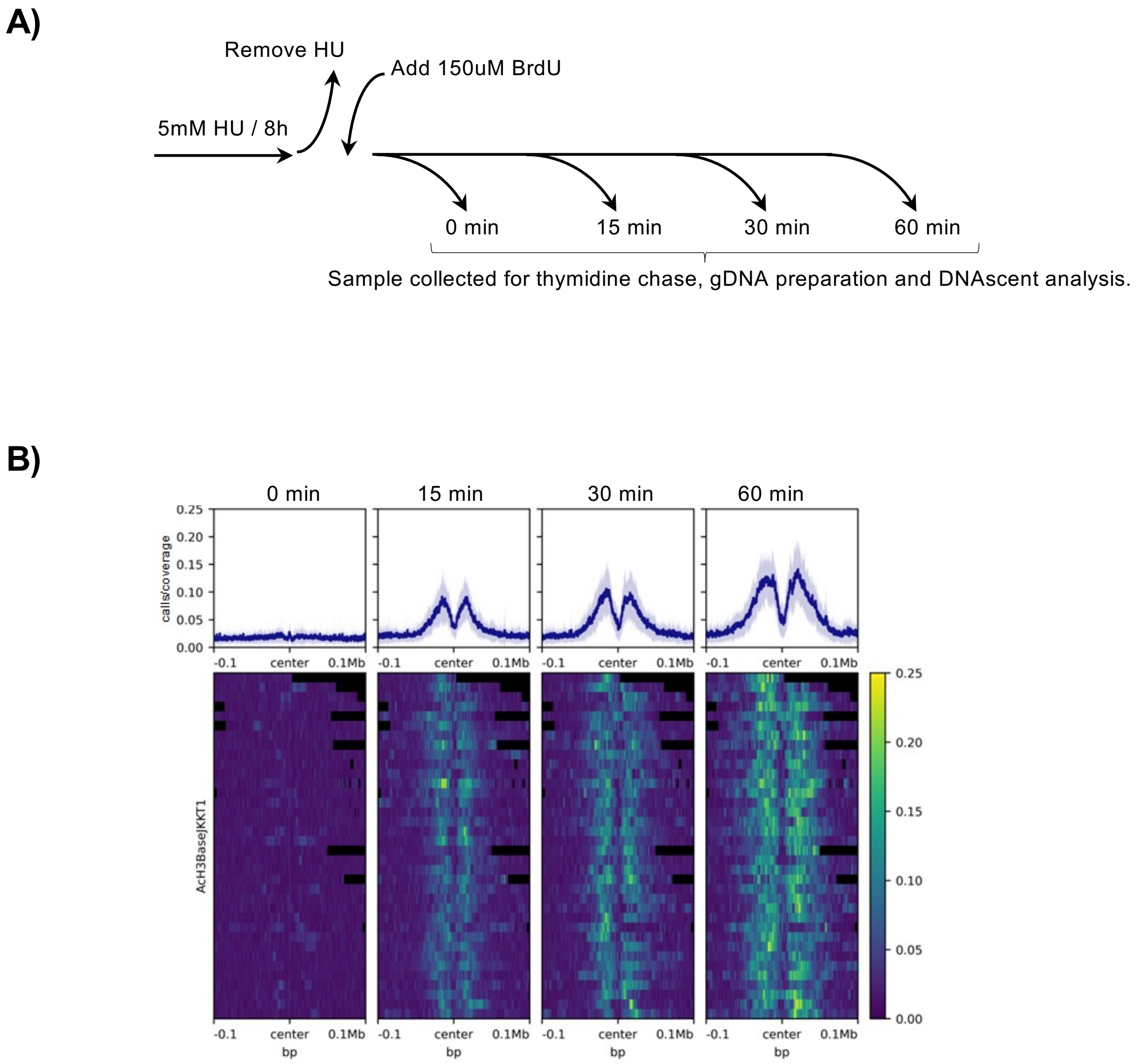

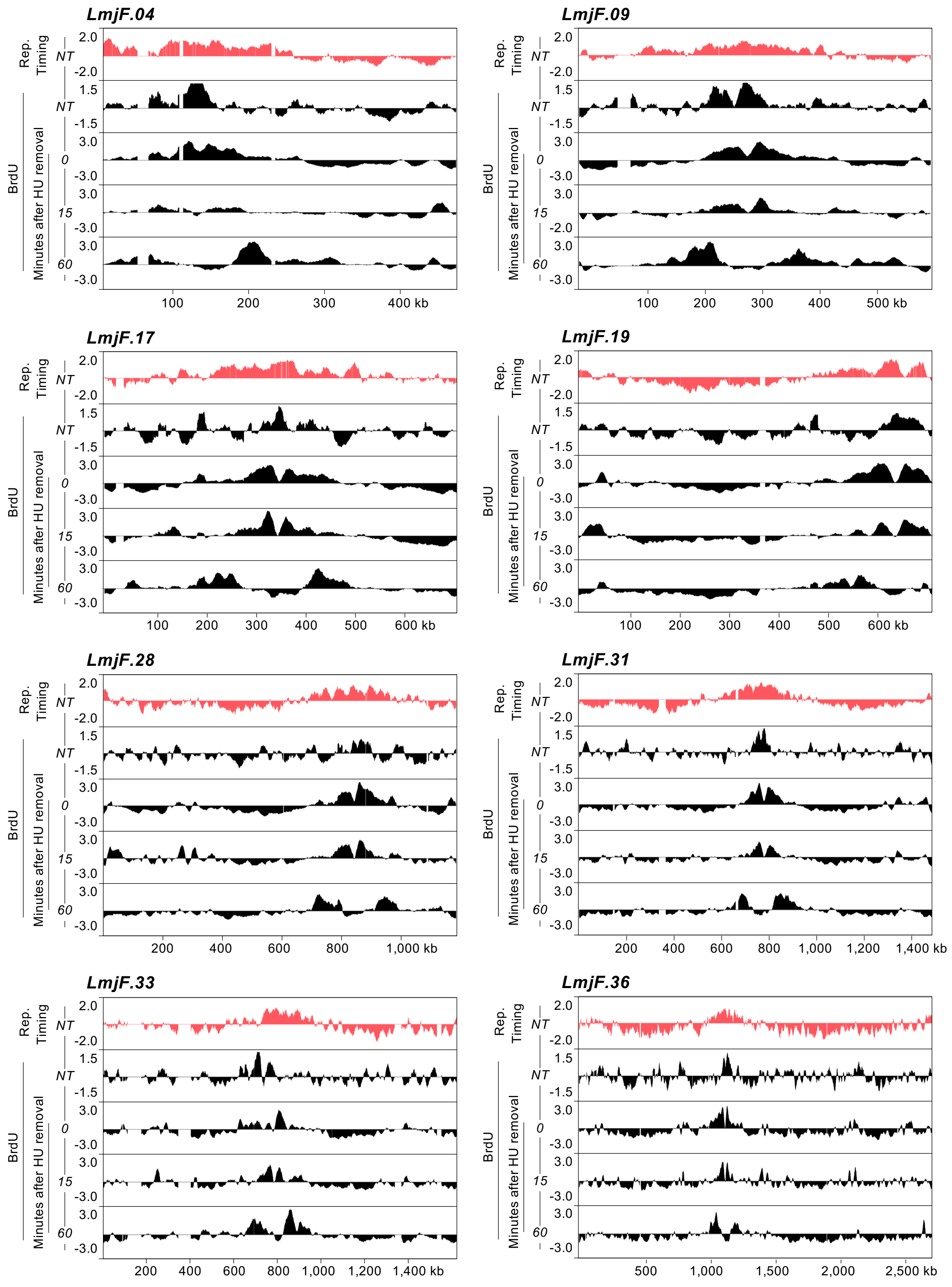

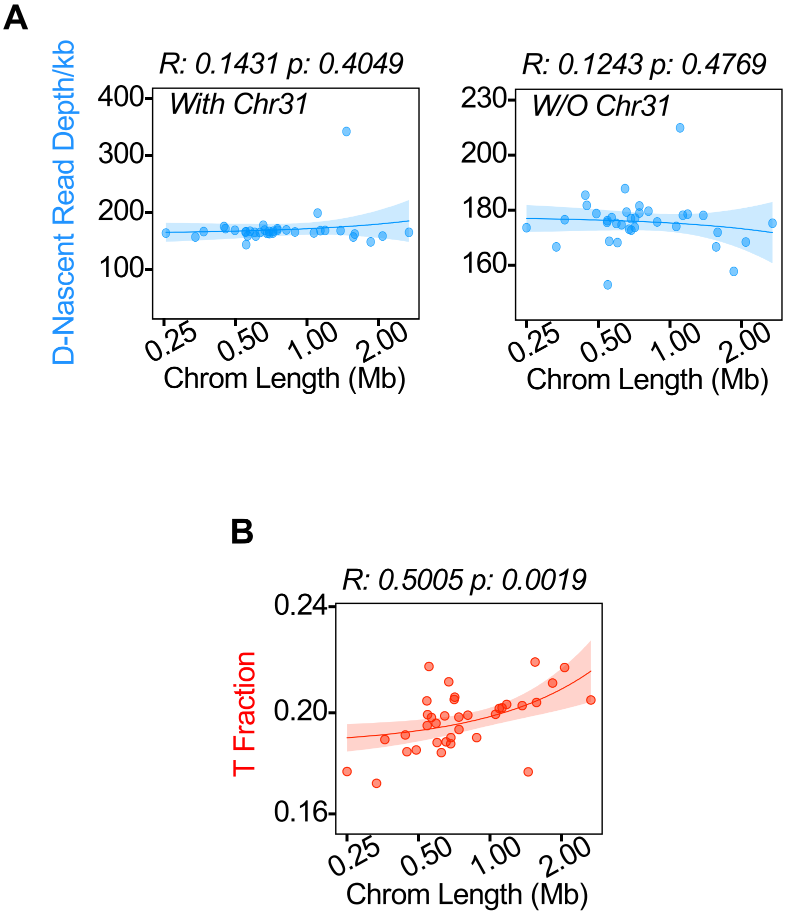

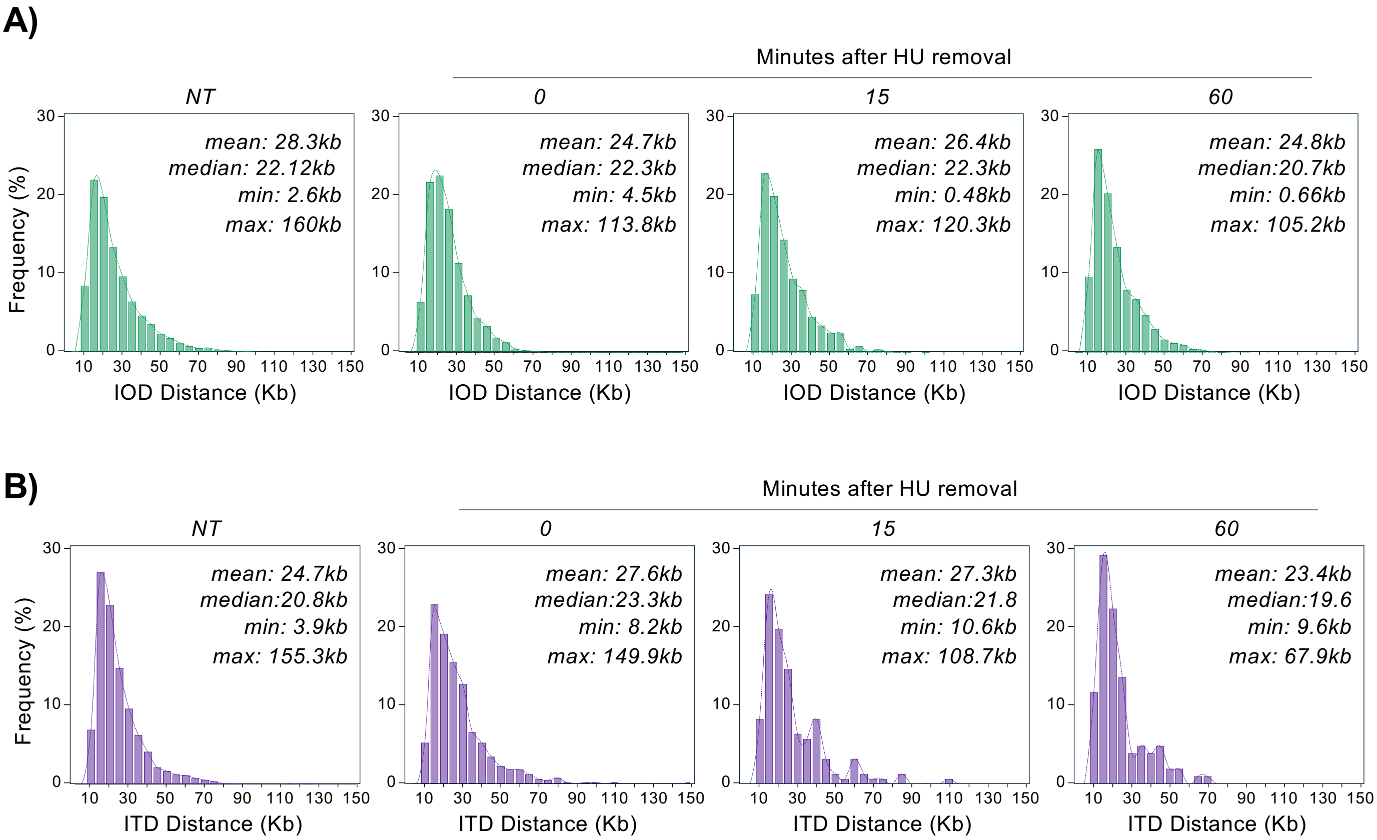

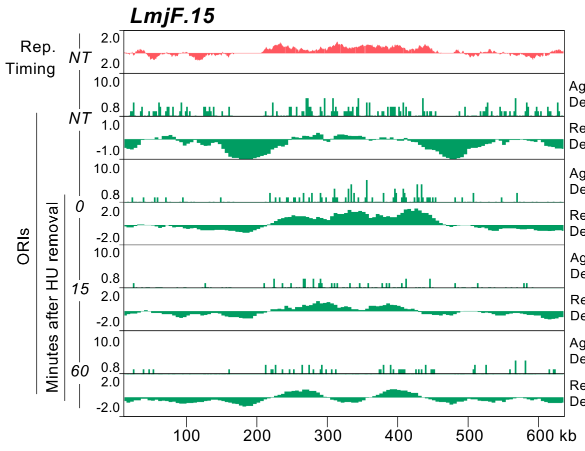

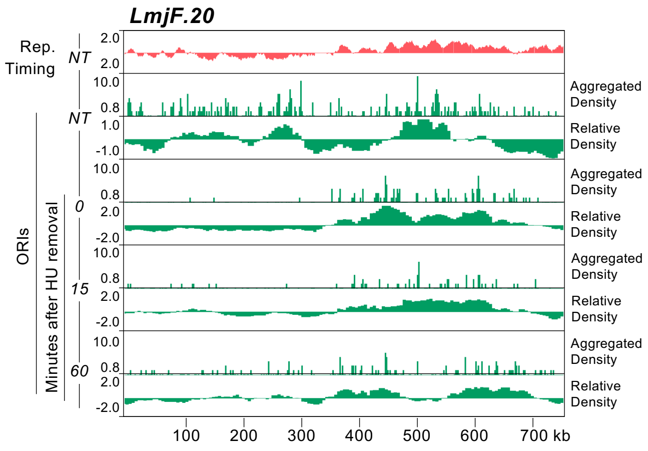

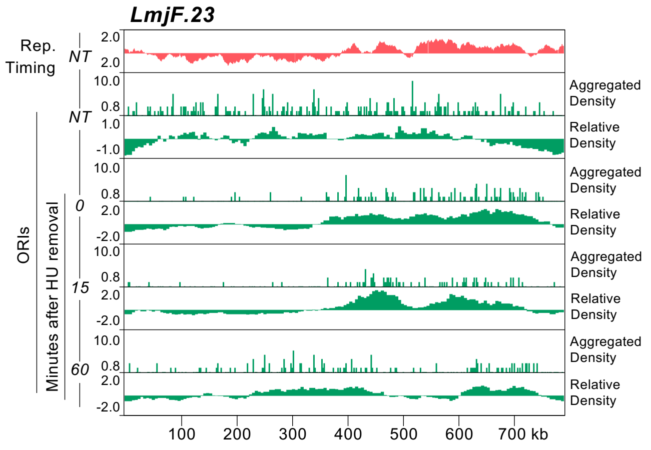

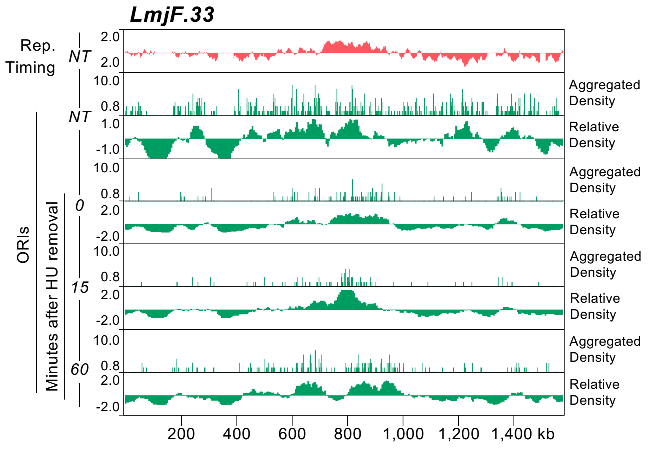

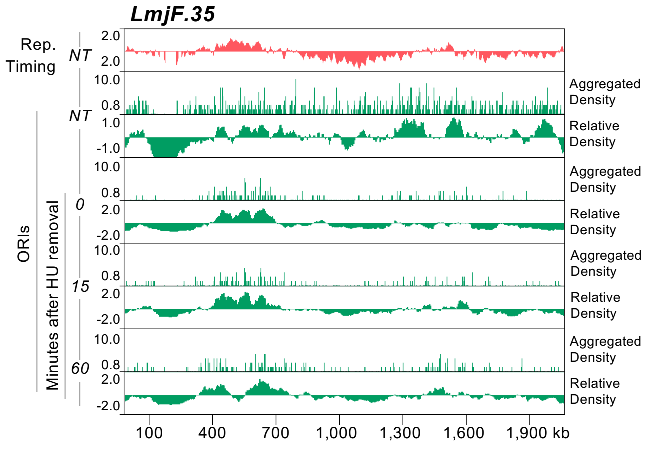

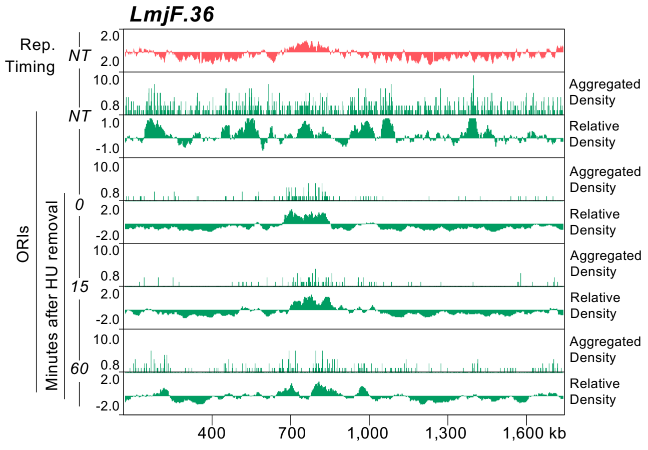

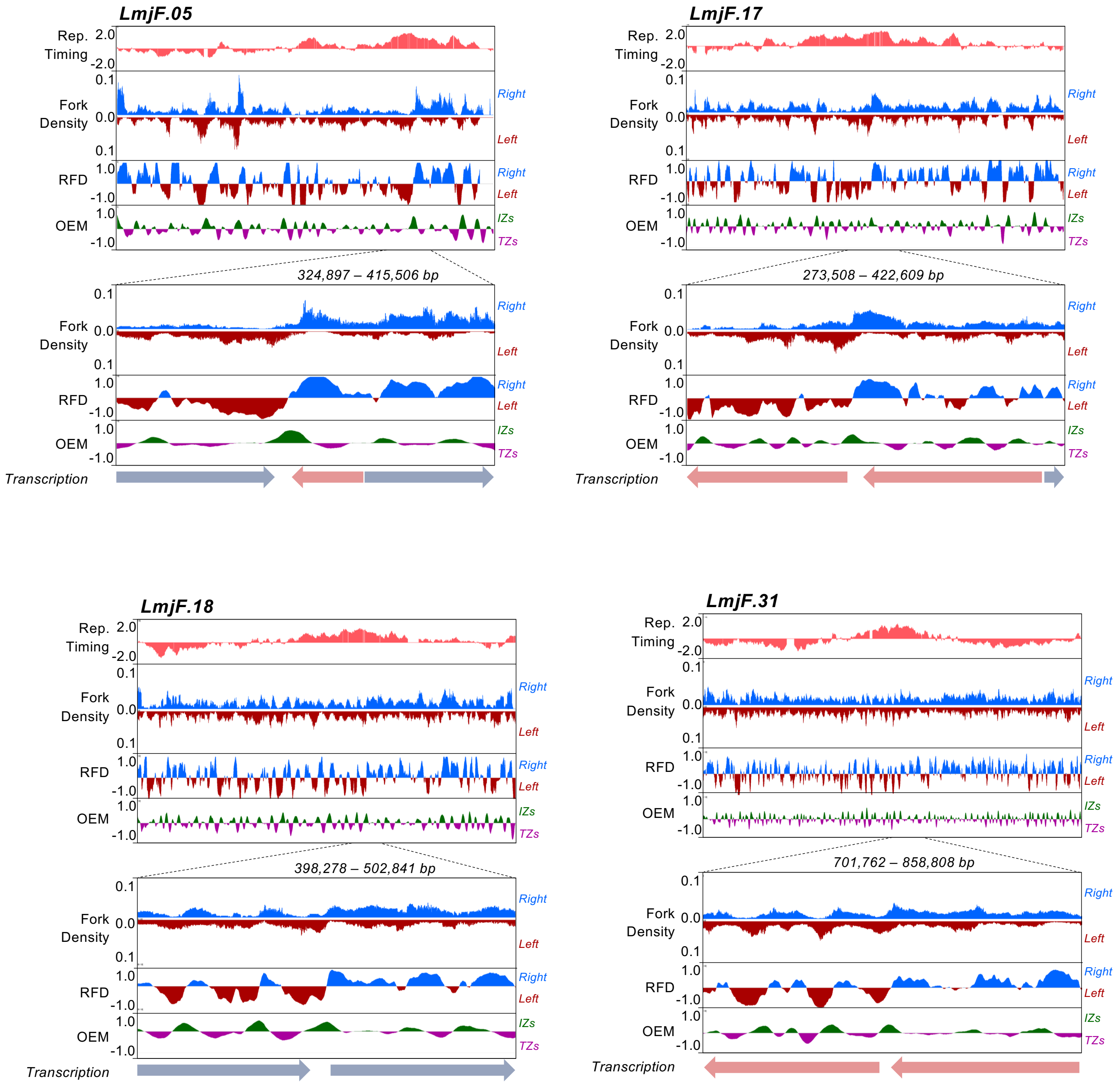

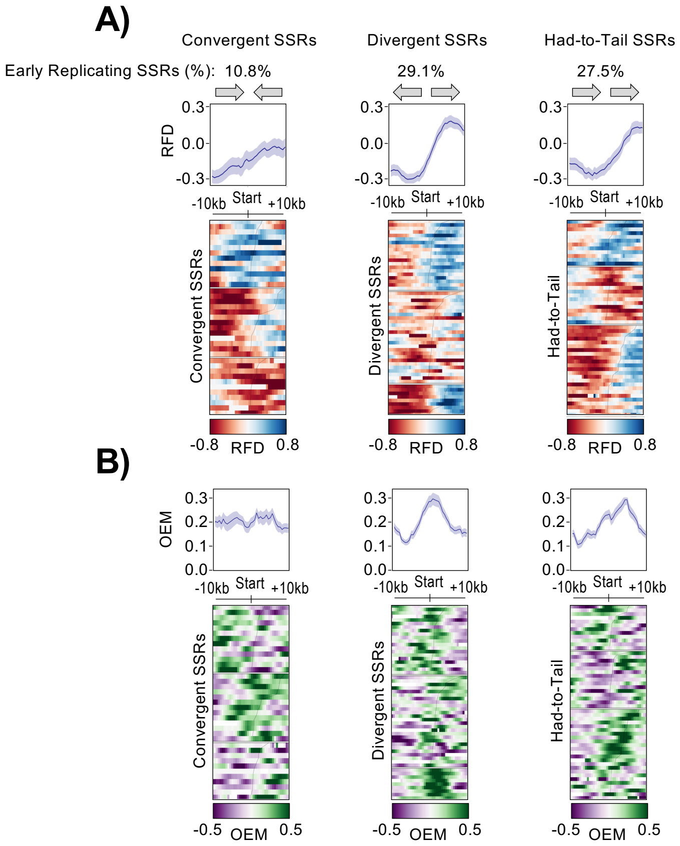

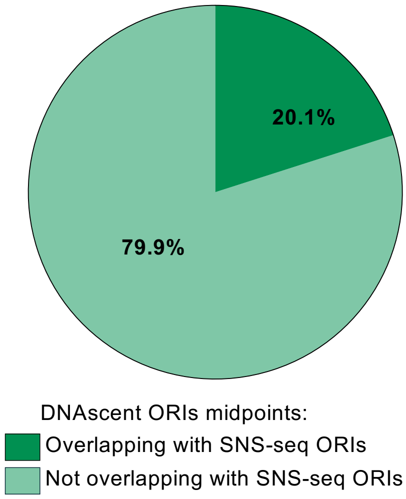

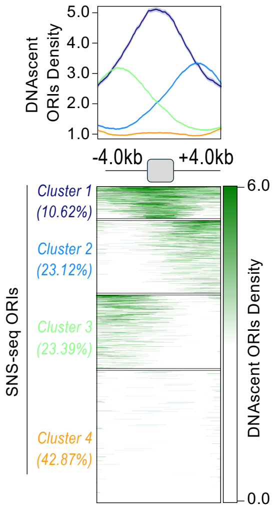
